# The Time and Place of European Admixture in Ashkenazi Jewish History

**DOI:** 10.1101/063099

**Authors:** James Xue, Todd Lencz, Ariel Darvasi, Itsik Pe'er, Shai Carmi

## Abstract

The Ashkenazi Jewish (AJ) population is important in medical genetics due to its high rate of Mendelian disorders and other unique genetic characteristics. Ashkenazi Jews have appeared in Europe in the 10^th^ century, and their ancestry is thought to involve an admixture of European (EU) and Middle-Eastern (ME) groups. However, both the time and place of admixture in Europe are obscure and subject to intense debate. Here, we attempt to characterize the Ashkenazi admixture history using a large Ashkenazi sample and careful application of new and existing methods. Our main approach is based on local ancestry inference, assigning each Ashkenazi genomic segment as EU or ME, and comparing allele frequencies across EU segments to those of different EU populations. The contribution of each EU source was also evaluated using *GLOBETROTTER* and analysis of IBD sharing. The time of admixture was inferred using multiple tools, relying on statistics such as the distributions of EU segment lengths and the total EU ancestry per chromosome and the correlation of ancestries along the chromosome. Our simulations demonstrated that distinguishing EU vs ME ancestry is subject to considerable noise at the single segment level, but nevertheless, conclusions could be drawn based on chromosome-wide statistics. The predominant source of EU ancestry in AJ was found to be Southern European (≈60-80%), with the rest being likely Eastern European. The inferred admixture time was ≈35 generations ago, but multiple lines of evidence suggests that it represents an average over two or more admixture events, pre-and post-dating the founder event experienced by AJ in late medieval times, with the pre-bottleneck admixture event bounded between 25-55 generations ago.

**Author Summary:** The Ashkenazi Jewish population has dwelt in Europe for much of its 1000-year existence. However, the ethnic and geographic origins of Ashkenazi Jews are controversial, due to the lack of reliable historical records. Previous genetic studies have exposed links to Middle-Eastern and European ancestries, but the history of admixture in Europe has not been studied in detail yet, partly due to technical difficulties in disentangling signals from multiple admixture events. Here, we address this challenge by presenting an in-depth analysis of the sources of European gene flow and the time of admixture events, using a wide spectrum of genetic methods, extensive simulations, and a number of new approaches. Specifically, to ensure minimal confounding by the Ashkenazi Middle-Eastern ancestry, we mask out genomic regions with Middle-Eastern ancestry, and investigate the lengths and geographic sources of the remaining regions. Our results suggest a model of at least two events of European admixture. One event slightly pre-dated a late medieval founder event and was likely from a Southern European source. Another event post-dated the founder event and was likely in Eastern Europe. These results, as well as the methods introduced, will be highly valuable for geneticists and other researchers interested in Ashkenazi Jewish origins and medical genetics.

## Introduction

Ashkenazi Jews (AJ), numbering approximately 10 million worldwide [1], are individuals of Jewish ancestry with a recent origin in Eastern Europe [2]. The first individuals to identify as Ashkenazi appeared in Northern France and the Rhineland (Germany) around the 10^th^ century [3]. Three centuries later, Ashkenazi communities emerged in Poland, due to migration from Western Europe and/or possibly from other sources. The Ashkenazi communities in Poland have grown rapidly, reaching millions by the 20^th^ century and wide geographic spread around Europe [2].

Due to the migratory nature of the Ashkenazi population and the relative scarcity of relevant historical records, the ethnic origins of present-day Ashkenazi Jews remain highly debated [2]. In such a setting, genetic variation provides crucial information. A number of recent studies have shown that Ashkenazi individuals have genetic ancestry intermediate between European and Middle-Eastern [4–8], consistent with the long-held theory of a Levantine origin followed by partial assimilation in Europe, and with the high observed genetic similarity to other Jewish communities. The estimated amount of accumulated European gene flow varied between studies, with the most recent ones, employing genome-wide data, converging to a contribution of about 50% to the AJ gene pool [4, 7, 9].

Despite these advances, very little is known about the identity of the European admixing population(s) or the time of the admixture events [2, 10], even though those are critical for our understanding of the origins of the early Ashkenazi Jews. Speculations abound due to the wide geographic dispersion of Jewish populations since medieval times [2], but only few historical records exists, underscoring the importance of genetic studies. Further complicating the picture is an Ashkenazi-specific founder event that has taken place about a millennium ago, as manifested by elevated frequencies of disease mutations [11, 12], reduced genetic diversity [13, 14], and abundance of long tracts of identity-by-descent [9, 15, 16]. Preliminary results from our recent studies [9, 17] were not decisive regarding the relative times of the European admixture and the founder event, calling for a more thorough investigation.

Some previous population genetic studies have attempted, often implicitly, to “localize” the Ashkenazi genomes to a single geographic region or source population [4–6, 18]. However, such approaches are confounded by the mixed European and Middle-Eastern Ashkenazi ancestry, which necessarily implies the existence of multiple sources. Here, we overcome this obstacle, following studies in other populations [19, 20], by performing a preliminary step of *local ancestry inference* (LAI), in which each locus in each Ashkenazi genome is assigned either a European or a Middle-Eastern ancestry. Following LAI, the source population of the European and Middle-Eastern “sub-genomes” can be determined independently, avoiding the “averaging” effect of treating the entire genome as a whole.

More specifically, we begin by testing the ability of available LAI software packages to correctly infer ancestries for simulated European/Middle-Eastern genomes. Proceeding with *RFMix*, we apply LAI to Ashkenazi SNP array data, and use a maximum likelihood approach to localize, separately, the European and Middle-Eastern sources. We show by simulations that our inference is robust to potential errors in the LAI. We also employ other methods based on allele frequency divergence between Ashkenazi Jews and other populations, although they turn out to be less informative. To estimate the time of admixture, we first use the lengths of European and Middle-Eastern tracts (calibrated by simulations) and the decay in ancestry correlations along the genome. We further introduce and apply a new method for dating admixture times based the genome-wide European or Middle-Eastern ancestry proportions. We integrate these results with an analysis of IBD sharing both within AJ and between AJ and other populations. Finally, compare our estimates to those produced by the *fineSTRUCTURE/GLOBETROTTER* suite [21–23]. Our results suggest that the European gene flow was predominantly Southern European (≈60-80%), with the remaining contribution either from Eastern or Western Europe. The time of admixture, under a model of a single event, is estimated to be around 30-45 generations ago. However, this admixture time is likely the average of at least two distinct events. Based on various lines of evidence, we propose that admixture with Southern Europeans pre-dated the late medieval founder event, whereas a more minor event in Eastern Europe was more recent.

## Results

### Data collection

SNP array data for Ashkenazi Jewish individuals was available from the schizophrenia study reported by Lencz et al., 2013 [24] (see also [25]). SNP arrays for European and Middle-Eastern populations were collected from a number of sources (*Table 1*). All genotypes were uniformly cleaned, merged, and phased (*Methods*), resulting in 2540 AJ, 543 European, and 293 Middle-Eastern genotyped at 252,358 SNPs. Note that while there are additional studies in these populations, we restricted ourselves to (publicly available) Illumina array data to guarantee a sufficient number of SNPs. We divided the European genomes into four regions: Iberia, North-Western Europe (henceforth Western Europe), Eastern Europe, and Southern Europe (Italy and Greece). The Middle-Eastern genomes were divided into Levant, Southern Middle-East, and Druze. See *Table 1* for further details and Figure S1 for a PCA plot supporting the partition into the indicated regions.

**Table 1.**
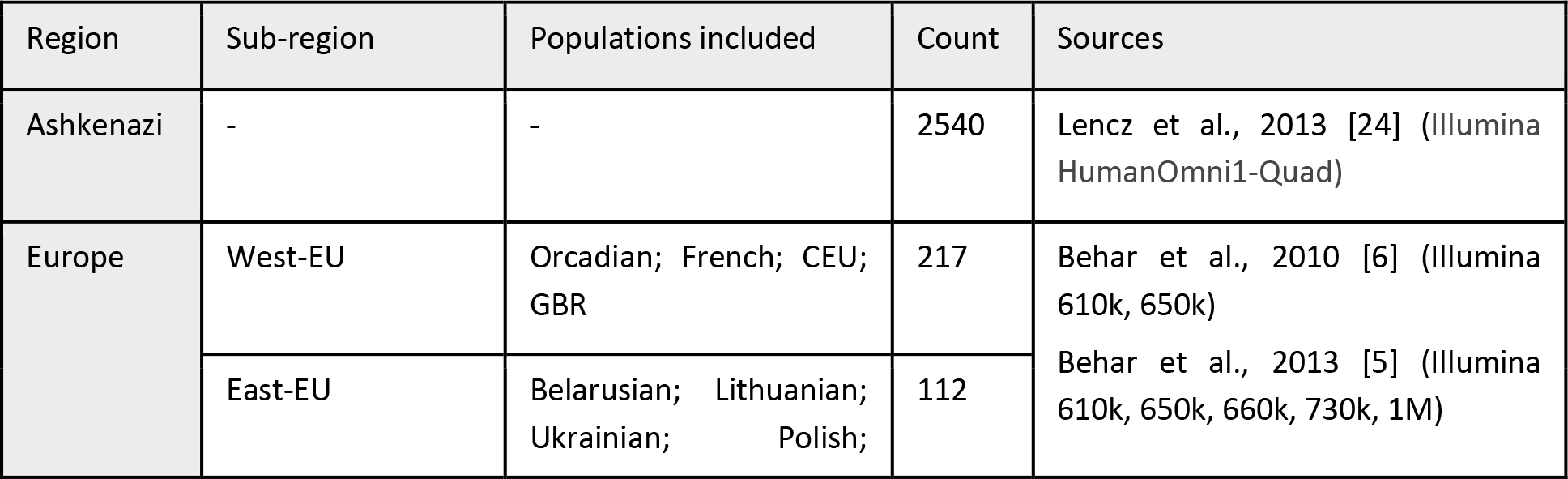
The populations and datasets used in our analysis.

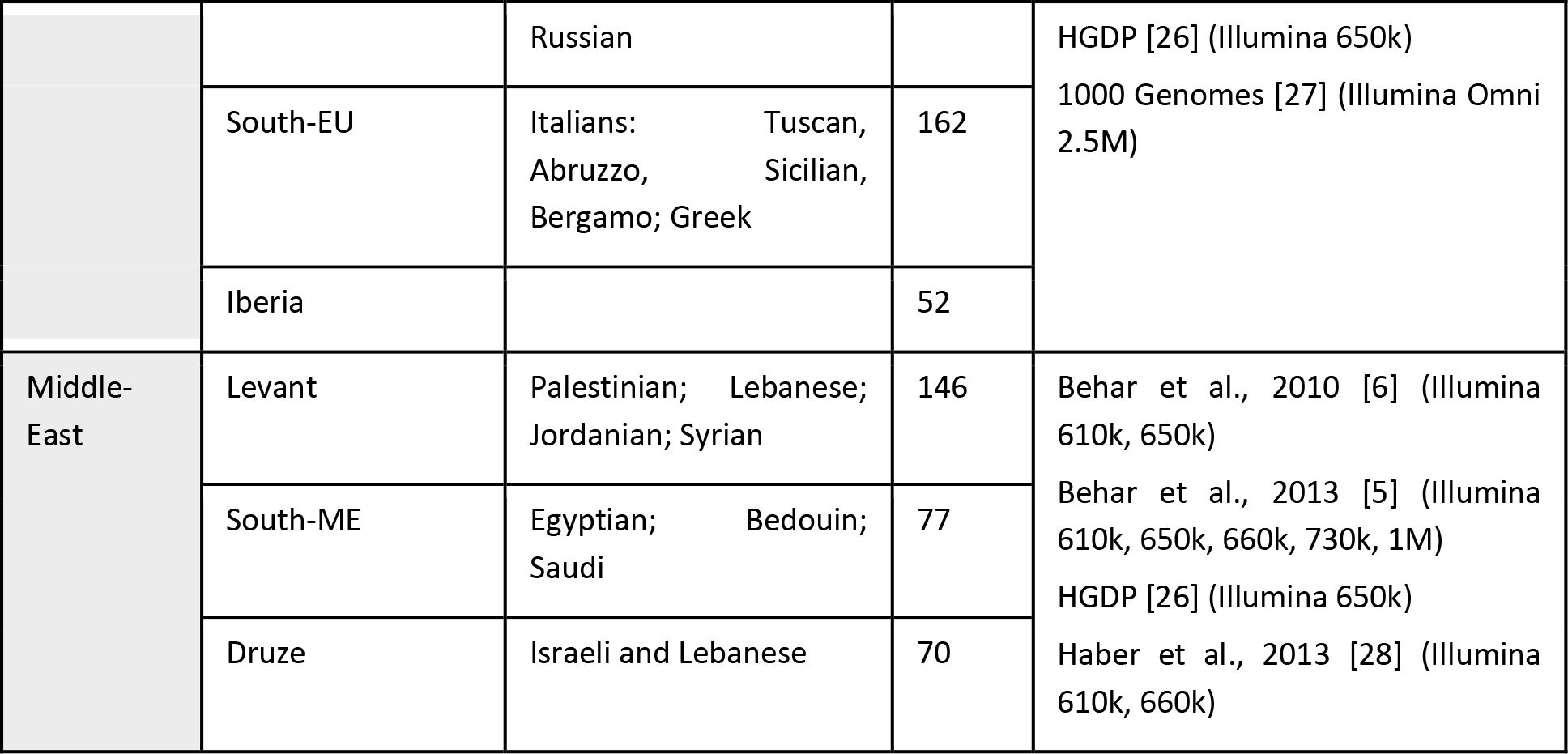

### Inferring the place of admixture using local ancestry inference

#### Calibration of the local ancestry inference method

In local ancestry inference (LAI), each region in the genome of each admixed individual is assigned an ancestry from one the reference panels. After evaluating the performance of LAI tools on admixture between closely related populations (Supplementary Text S1), we selected *RFMix* [29], which is based on a random forest classifier for each genomic window and smoothing by a hidden Markov model. When running *RFMix*, we did not iterate over the inference process using the already classified individuals (the Expectation-Maximization step), as we found that accuracy did not improve (*Methods*) and we wanted to avoid bias due to the widespread haplotype sharing typical to the AJ population. We also did not filter SNPs by the quality of their local ancestry assignment, as we found that such filtering substantially biases downstream inferences (Supplementary Text S1). Finally, we downsampled the reference panels to balance the sizes of the European and Middle-Eastern sample sizes, as well as balanced the number of genomes from each European region (*Methods*).

Running RFMix on the AJ genomes with our European and Middle-Eastern reference panels and summing up the lengths of all tracts assigned to each ancestry, the genome-wide ancestry was ≈53% EU and ≈47% ME, consistent with an *ADMIXTURE* analysis (*Methods*) and our previous estimate based on a smaller sequencing panel [9]. Our simulations suggested that the accuracy of LAI for an EU-ME admixed population is only around ≈70-80%, much lower than the near-perfect accuracy observed for cross-continental admixture (e.g., [29–33]). Even so, the local ancestry assignment is still far from being random, and therefore, with proper accounting for errors (below), it is informative on the place and time of admixture events.

#### Geographic localization of the EU component of the AJ genomes

Following the deconvolution of segments of EU and ME ancestries, we focused on the regional ancestry of the European segments. We initially followed refs. [19, 20] and attempted to apply *PCAMask* to the EU subset of the AJ genomes. However, *PCAMask’*s results were inconsistent across runs and parameter values (see Supplementary Text S2 and [34]). We therefore developed a simple naïve Bayes approach. We first thinned the SNPs to assure linkage equilibrium between the remaining SNPs. We then computed the allele frequencies of the SNPs in the four geographical regions: Southern EU, Western EU, Eastern EU, and Iberia. Then, for each haploid chromosome, we computed the log-likelihood of the European assigned part of the chromosome to come from each of the four regions as a simple product of its allele frequencies, normalized by the number of European classified SNPs at each chromosome.

Initial inspection of the results revealed that Iberia had consistently lower likelihood than the other regions. We therefore removed the Iberian genomes, and since the Iberia panel was the smallest and sample sizes had to be balanced across regions, this enabled us to increase the sample size for the other regions (*Methods*). To determine whether the true ancestry could theoretically be recovered given a single European source, we generated simulated chromosomes using genomes not included in the *RFMix* reference panel. Each simulated chromosome was a mosaic of segments from Middle-Eastern and European genomes, and segment lengths were exponentially distributed, according to the expected parameters of a symmetric admixture event taking place 30 generations ago (*Methods*). In each simulation experiment, the identity of the European source region was varied, and the log-likelihood was averaged over all chromosomes. Running the same pipeline as for the real data, we were able to correctly identify the source in all three cases (*Figure 1*). This result indicated that localization of the European source is feasible, despite noise and biases in local ancestry inference between closely related population such as Middle-Easterners and Europeans.

**Figure 1.**
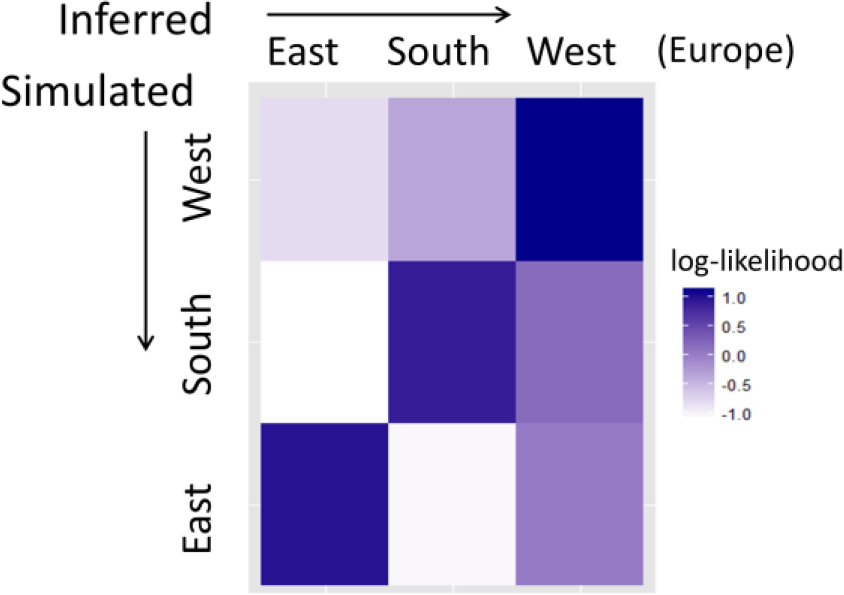
Simulation results for our localization pipeline. In each row, admixed genomes were simulated with sources from the Levant (50%) and one European region (50%). Columns correspond to the inferred log-likelihood of each potential source.

For the AJ data, we found that Southern Europe was the most likely source for the EU component of the largest proportion of the AJ chromosomes. Specifically, 43.2% of the AJ chromosomes had Southern EU as their most likely source, 35.4% had Western EU and 18.8% had Eastern EU (the proportions do not precisely sum to 1, as we allowed, for control, classification as Middle Eastern). Therefore, Southern Europe is the dominant source of gene flow into AJ. Nevertheless, we did not yet quantify the magnitude of the Southern EU component and of other, minor sources.

For the Middle-Eastern source, we observed that in simulations of admixed genomes, the Middle-Eastern regional source could also be recovered by running the same localization pipeline (not shown). Applying this pipeline to the AJ genomes, we identified Levant as the most likely ME source.

#### The magnitude and identity of the minor European components

To estimate the contribution of each subcontinental European region, we performed 4-way admixture simulations between individuals of Levantine, Southern European, Eastern European, and Western European origin. In these simulations, we fixed the Levant admixture proportion to 50% and varied the proportions of different European regions. We then used a grid-search to find the ancestry proportions that best fit the observed fraction of AJ chromosomes classified as descending from each ancestry, as described in the previous section. The simulation results (*Figure 2*) suggested that the European component of the AJ cohort is composed of 34% Southern EU, 8% Western EU, and 8% Eastern EU ancestries. This analysis thus suggests that roughly 70% of EU ancestry in AJ is Southern European. Using bootstrapping, the 95% confidence interval of the Southern EU ancestry was [33,35]% and that of Eastern EU was [8,9]%. Note that while the mean likelihood of Southern EU was only very slightly higher than Eastern/Western EU (not shown), our simulations clearly showed that this observation is consistent with a predominant Southern EU source. We hypothesize that this is due to ME segments being more distinguishable from Northern segments than from the more closely related Southern EU ones. This differential detection then leads to an enrichment of Northern EU ancestry among the inferred EU segments, and underscores the importance of our simulations.

**Figure 2.**
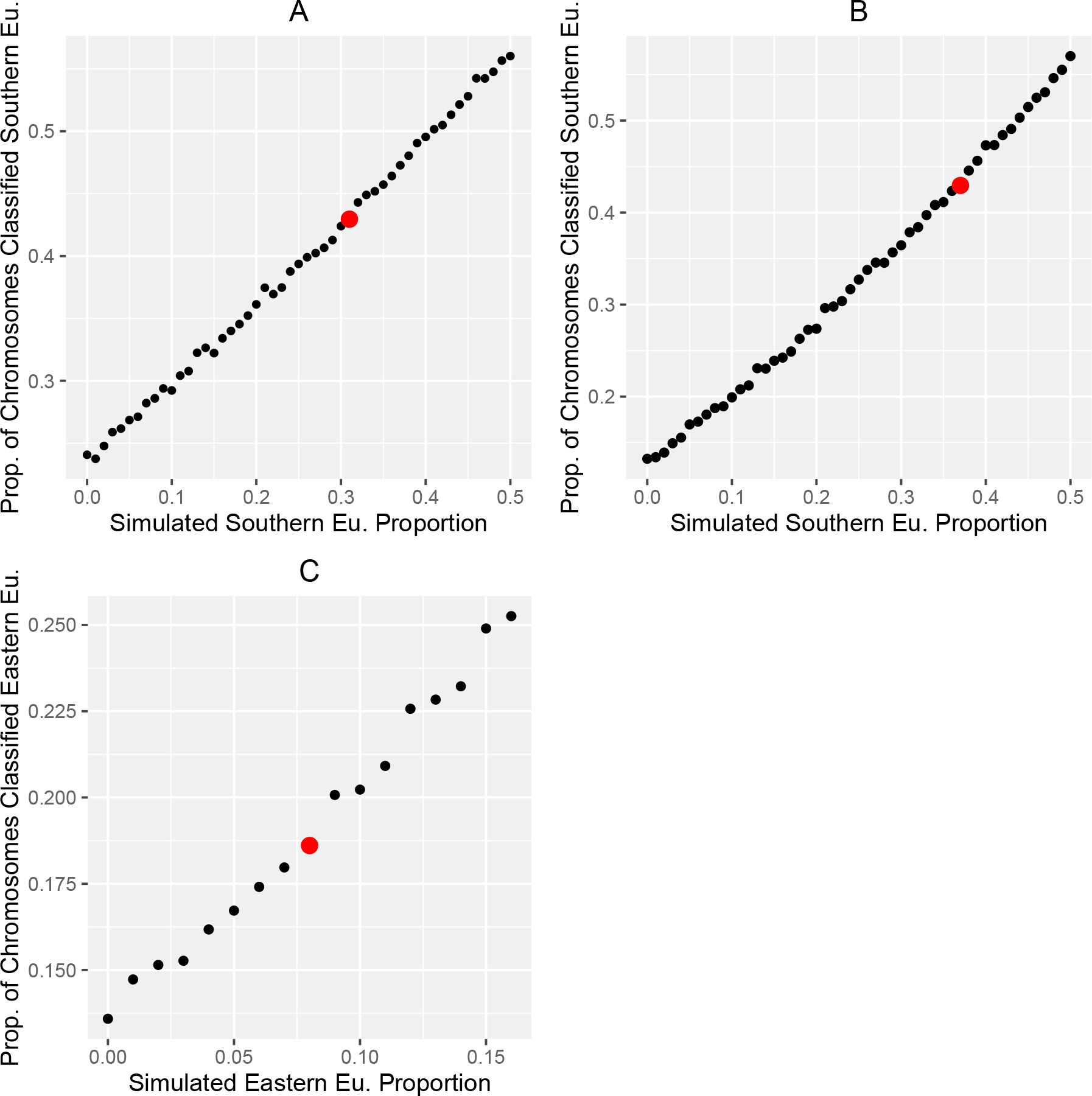
Inference of the proportion of Ashkenazi ancestry deriving from each European region. We simulated admixed chromosomes with European and Middle-Eastern ancestries, where the ME ancestry was fixed to the Levant region and to 50% of the overall ancestry. We then varied the sources of the remaining European ancestry to determine which ancestry proportions most closely match the AJ data. In (A), the simulated EU component was Southern and Western EU. For each given proportion of Southern EU ancestry, we used our LAI-based pipeline to compute the proportion of chromosomes naïve-Bayes-classified as Southern European. The best match to the proportion of thus classified chromosomes observed in the real AJ data (red dot) was found when the true simulated Southern EU ancestry was 31% of the total. In (B), the same simulation procedure was repeated, except that the simulated EU components were of Southern and Eastern EU ancestry. The inferred proportion of Southern EU ancestry in AJ is now 37%. (C) We fixed the Southern EU contribution to 34%, the average of its estimates from (A) and (B), and varied the remaining 16% between Western and Eastern EU. The simulations suggest that the closest match to the real results is at roughly equal (8%) Western EU and Eastern EU ancestry proportions. Bootstrapping was used to obtain confidence intervals by resampling AJ individuals 1000 times with replacement;
to obtain the simulated value matching each bootstrap iteration, we used linear regression in the region near the real AJ value.

### Inferring the time of admixture using local ancestry inference

#### Mean segment length

Consider a model of a “pulse” admixture between two populations, *t* generations ago, with respective proportions *q*:(1-*q*). The mean length (in Morgans) of segments coming from the second source is 1/(*qt*) [35]. In the case of AJ, where the source populations are EU and ME, we estimated *q* above as ≈53%. Therefore, the mean ME (or EU) segment length is expected to be informative on the time of admixture *t*. The mean ME segment length was ≈14cM; however, we noticed that in simulations, the *RFMix*-inferred segment lengths were significantly overestimated. To correct for that, we used simulations to find the admixture time that yielded *RFMix*-inferred segment lengths that best matched the real AJ data. In the simulations, we fixed the ancestry proportions to the ones inferred above for AJ (50% ME, 34% Southern EU, 8% Western EU, and 8% Eastern EU), and varied the admixture time. We then plotted the RFMix-inferred ME segment length vs the simulated segment lengths (*Figure 3*). The simulated mean segment length that corresponds to the observed AJ value was around 6.6cM, which implies an admixture time of ≈29 generations ago (95% confidence intervals: [27,30] generations).

**Figure 3.**
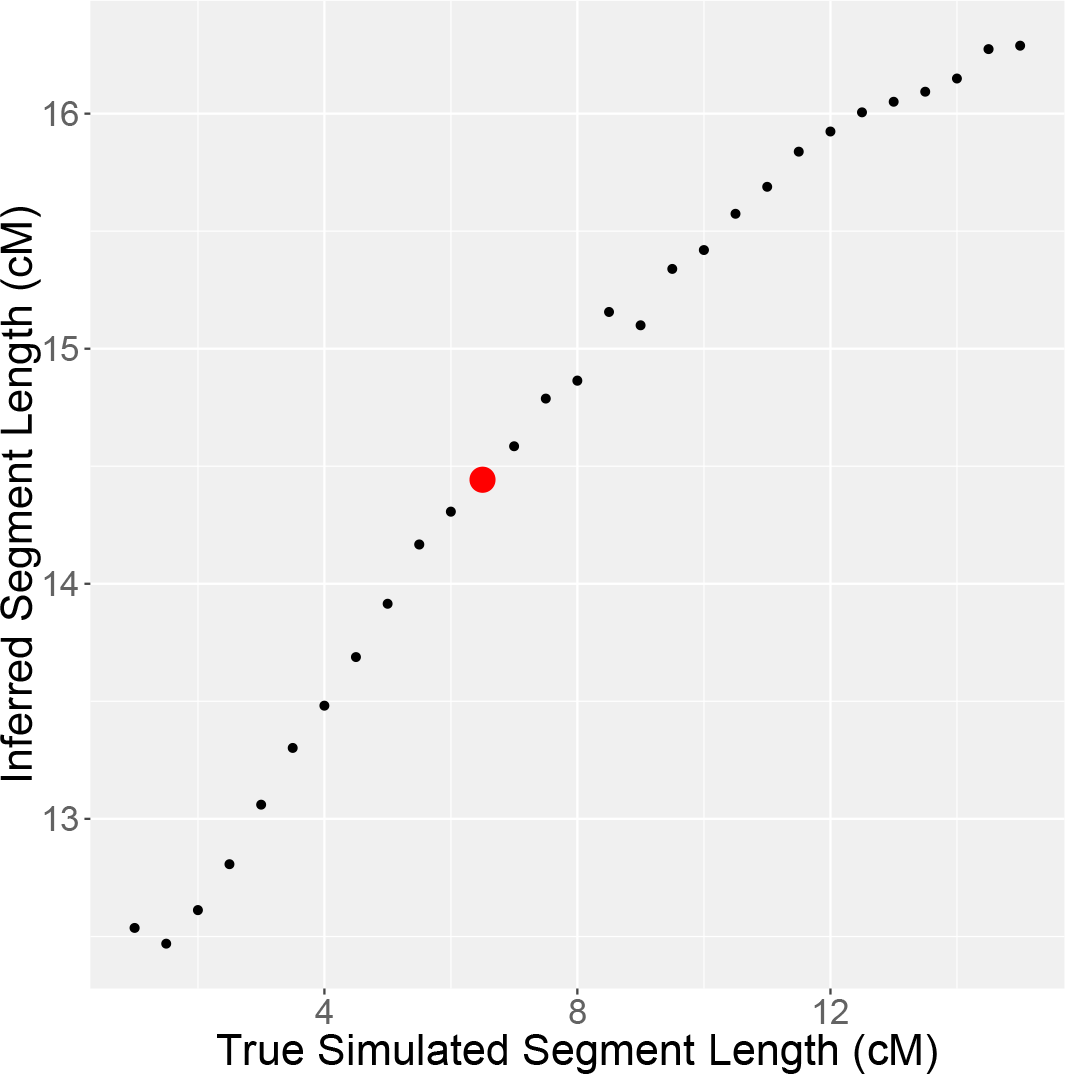
Inferring the AJ admixture time using the lengths of admixture segments. The mean length of *RFMix*-inferred Middle-Eastern segments is plotted vs the mean simulated length, which is inversely correlated to the simulated admixture time. The red dot corresponds to the observed mean segment length in the real AJ data. Confidence intervals were computed as in **Figure 2**.

#### Chromosome-wide ancestry proportions

Beyond mean segment lengths, the proportion of ancestry (per chromosome) that descends from each ancestral population is also informative on the time of admixture [36, 37], since the longer the time since admixture, the smaller its variance [35]. While ancestry proportions contain less information than segment lengths, they are expected to be more robust to misidentification of the segment boundaries. Building on models from refs. [35, 38, 39], we derived a new analytical expression for the distribution of ancestry proportions (for either phased or unphased data) given the initial admixture proportions and admixture time (see *Methods* for details). Given observed ancestry proportions, we then obtained a maximum likelihood estimate of the admixture time and the initial proportions. For admixture between highly diverged populations, the method is expected to work well for intermediate admixture times (say, 10 < *t* < 200s generations [40]), as we demonstrated using simulations (Figure S2).

To apply our method to the AJ admixture, we used the LAI results and summed up the lengths of European vs Middle-Eastern segments. While we could have estimated the ancestry proportions directly using tools like *ADMIXTURE* [41], experiments with simulated data demonstrated that for EU/ME admixture, LAI is much more accurate even for the chromosome-wide ancestry proportions (see Discussion). However, our simulations showed that even using LAI, for EU/ME admixture, the correlation between true and inferred ancestry proportions is only *r^2^* ≈ 0.11 (Figure S3). Therefore, the results from an application of our method on the AJ data (EU ancestry *q* = 0.55 and admixture time *t* = 22 generations) should be considered only as an order of magnitude estimate.

To correct for the distortion of the distribution caused by local ancestry inference, we again used EU/ME admixture simulations. We found that the best fit to the AJ data using a 4-way model (Middle-Eastern, Southern EU, Eastern EU, and Western EU with proportions 50:35:12:3 (%), respectively) was obtained with admixture time of 35 generations ago (*Figure 4*), close to the time inferred above using the mean segment lengths. This time is also consistent with the estimates from *Alder* and *GLOBETROTTER* described below.

**Figure 4.**
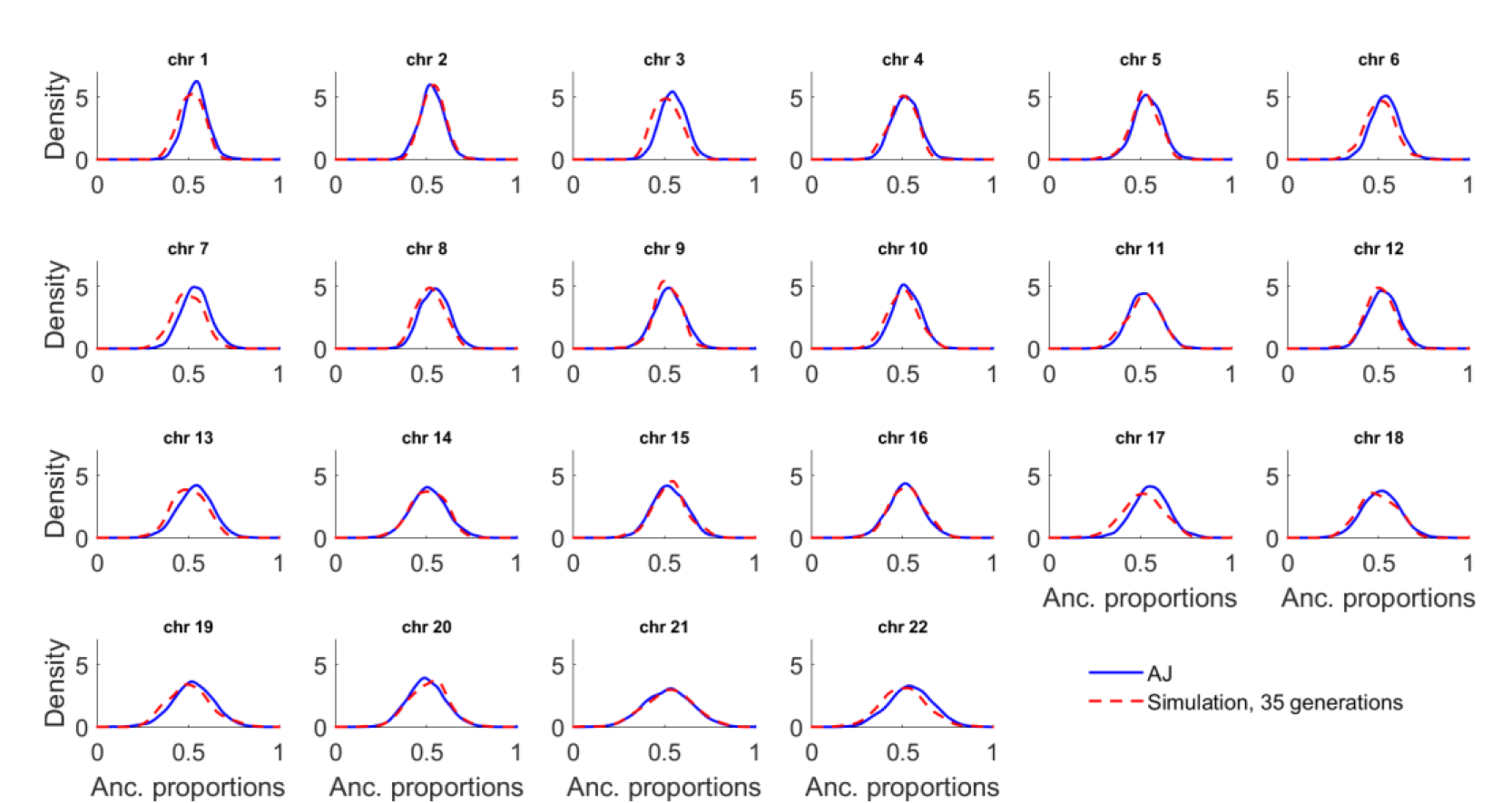
The density of ancestry proportions in AJ and in simulations. The ancestry proportions in AJ were computed using LAI (*RFMix*). Simulations are of 200 genomes with a history of an admixture pulse 35 generations ago between Middle-Eastern, Southern EU, Eastern EU, and Western EU populations. The density was estimated using a normal kernel.

The assumption of pulse admixture, however, might be unrealistic, in particular that we have identified more than two likely ancestral sources. In Supplementary Text S4, we analytically derived the distribution of segment lengths and ancestry proportions for a double admixture model, where the initial admixture event was followed by a second contribution from one of the sources. However, we empirically observed that the ancestry proportions from this model can often be fitted excellently by pulse admixture. Given this and the considerable noise introduced by LAI, directly estimating the parameters of multiple admixture events is unlikely to be reliable.

To overcome this problem, we first note that the inferred single admixture time, even if estimated using a simplified model, still imposes constraints on the admixture times and proportions in a double admixture model (*Methods*). Additionally, we notice that the estimated admixture time (≈30-35 generations) is very close to the estimated time of the AJ bottleneck event [9, 16]. If indeed multiple distinct admixture events have taken place, they must have necessarily happened on either side of the bottleneck, and thus leave different traces when examining the ancestry of genomic segments with ancestry at the bottleneck. We apply these insights in the following sections.

### Ancestry of identical-by-descent (IBD) segments

A number of recent studies have shown that sharing of identical-by-descent (IBD) segments is abundant in the AJ population, and is likely due to a severe bottleneck taking place around 30 generations ago [4, 7, 9, 15, 16]. An open question is the relative timing of the bottleneck and the European gene flow, with our current and past [9] point estimates dating the gene flow at around or slightly earlier than the bottleneck. Given that most long segments (e.g., with length >3cM and <7cM) coalesce around the time of the bottleneck, we contrast two hypothesis, as follows. If admixture completely predated the bottleneck, then IBD segments should have the same EU/ME ancestry proportions as observed genome-wide. If, on the other hand, gene flow from one source population entered AJ long after the bottleneck, then the ancestry of the IBD segments should be predominantly from the other source population (e.g., see [42–44]). Elevated ME ancestry of IBD segments would thus indicate European gene flow both before and after the bottleneck. Further, IBD segments shared between AJ and other populations could shed light on the geographic origin of each admixture event.

We detected IBD segments shared within AJ individuals using *Germline* [45] and *Haploscore* [46] (*Methods*). We then computed the total amount of genetic material in IBD segments associated with each diploid ancestry; namely, each segment was assigned as having either homozygous EU ancestry, homozygous ME ancestry, or heterozygous ancestry. Clearly, errors in IBD segment detection and local ancestry inference could severely bias the conclusions of our analysis. Fortunately, we could naturally and completely account for these by using the observed number of IBD segments shared between individuals labeled homozygous ME and homozygous EU, since the number of such segments is a direct measure of the noise level (*Methods*).

Our results demonstrated an over-representation of Middle-Eastern IBD segments, consistent with two waves of gene flow. We then estimated (*Methods*) the European fraction of the AJ ancestry at the bottleneck at 42%, less than the 53% observed genome-wide. The contribution of *post-bottleneck* European gene flow required to explain these figures is 19% of the AJ gene pool (*Methods*). Eliminating particularly long segments (>7cM; as those may derive from ancestors even more recent than the bottleneck), increased slightly the inferred magnitude of post-bottleneck gene flow to 22%, or 23% when considering only segments <4cM.

Given a history of multiple admixture events, a natural question is the geographic source of each. According to the documented AJ migration history, it is attractive to speculate that the Southern-European gene flow was pre-bottleneck and that the Western/Eastern European contribution came later. Indeed, we note that the estimated proportion of ≈20% post-bottleneck replacement is close to our above estimate of ≈16% EU gene flow from sources other than Southern-EU as well as to *TreeMiXs* and *Globetrotter’*s results below (and perhaps also with our previous estimate of ≈15% EU ancestry based on AJ and Western European (CEU) data alone [17]). To test this hypothesis, we considered the European ancestry of IBD segments longer than 15cM, which are highly unlikely to predate the bottleneck. Compared to the genome-wide results, the proportion of AJ individuals (with all regions masked but the >15cM IBD segments) inferred by our geographic localization pipeline (applied to entire individuals) to be most likely Southern European decreased by 14.8% points, with the proportion of AJ individuals inferred to be most likely Eastern and Western European increasing by 10.2 and 4.5% points, respectively. [As a control, when we considered AJ individuals reduced to IBD segments of *any* length, there was no noticeable difference from the genome-wide results.]

Finally, we considered IBD segments shared between AJ and other populations (*Figure 5*), and observed that the number of segments shared between AJ and Eastern European was ≈6-fold higher than shared between AJ and Southern Europeans (consistent with [5]), with this ratio increasing to ≈60-fold for segments of more recent origin (length >7cM). Further, the number of segments shared with Eastern Europeans was ≈2-fold higher than with Western Europeans or the people of Iberia (P=5·10^−3^ for the difference, using permutations of EU regional labels), pointing to Eastern Europe as the predominant source of the recent gene flow. We note though that IBD sharing between AJ and European individuals is a very rare event: the mean number of segments shared between AJ and Eastern Europeans is ≈0.04.

**Figure 5.**
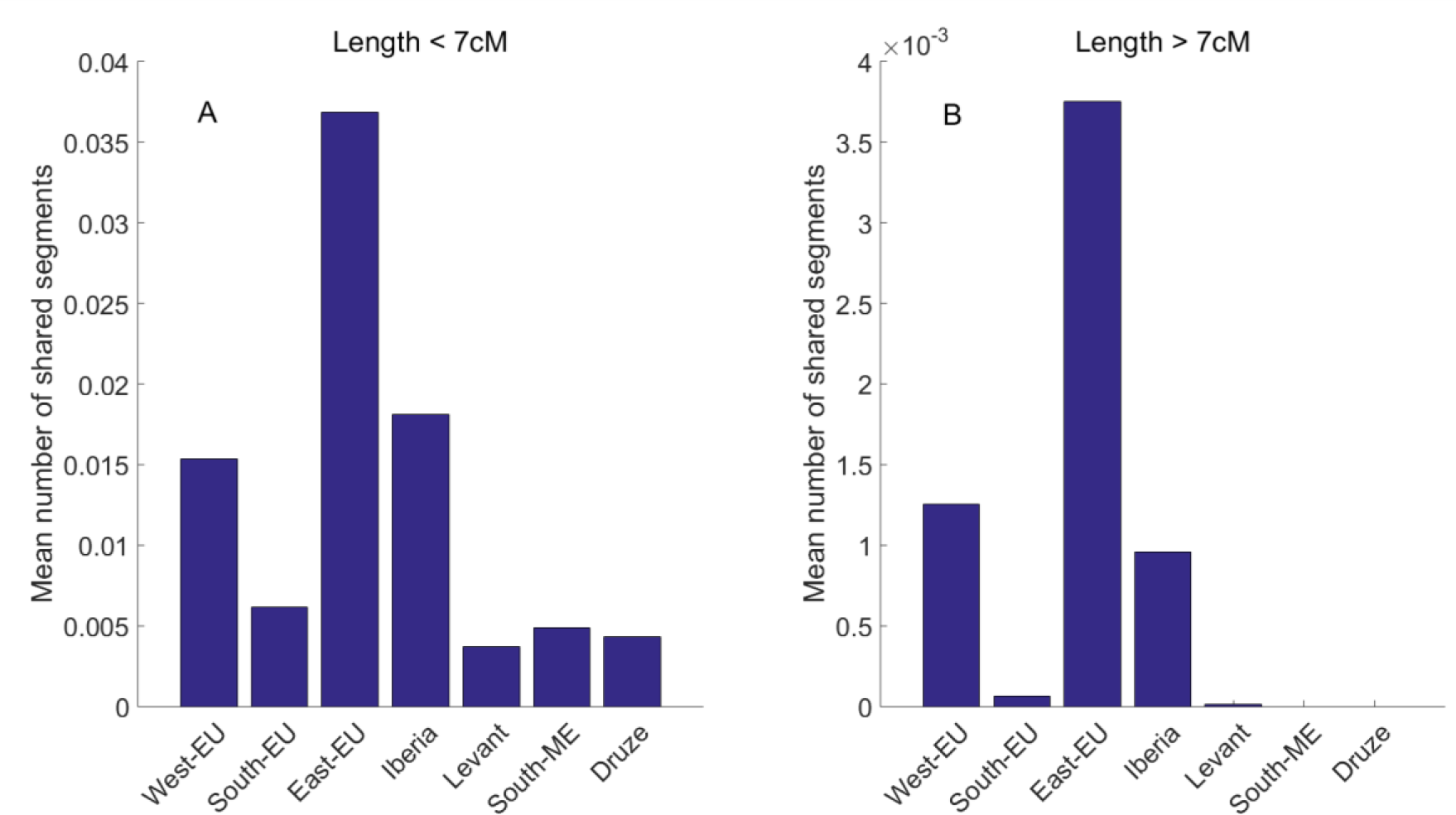
The number of IBD segments shared between Ashkenazi Jews (AJ) and other groups of populations. IBD segments were detected by *Germline* and *Haploscore* as explained in *Methods*. The population groups are as in **Table 1**. Note the different scale of panels (A) and (B) (segments of length between [3,7]cM and >7cM, respectively), and that sharing between AJ and either Southern Europeans or Middle Easterners completely vanishes for the longer (more recent) segments, indicating a relatively older divergence/gene flow. Also note that while sharing with Eastern Europeans is high compared to other groups, it is nevertheless a relatively rare event (≈0.04 segments per pair), in particular compared to sharing within AJ (≈3.4 segments per pair, almost 100-fold higher).

### Inferring the time and source of gene flow using additional methods

#### Decay of admixture linkage disequilibrium (Alder), f3/f4 statistics, and tree structure (Treemix)

Refs. [47–49] have shown that linkage disequilibrium (LD) in an admixed chromosome, weighted properly, decays exponentially with the genetic distance, and the *Alder* package was implemented to infer the admixture time and the ancestral sources. The admixture time inferred by *Alder* for AJ is broadly consistent with the LAI-based results, at 30-40 generations ago (Table 2; the P-value for admixture was significant under all tests).

**Table 2.**
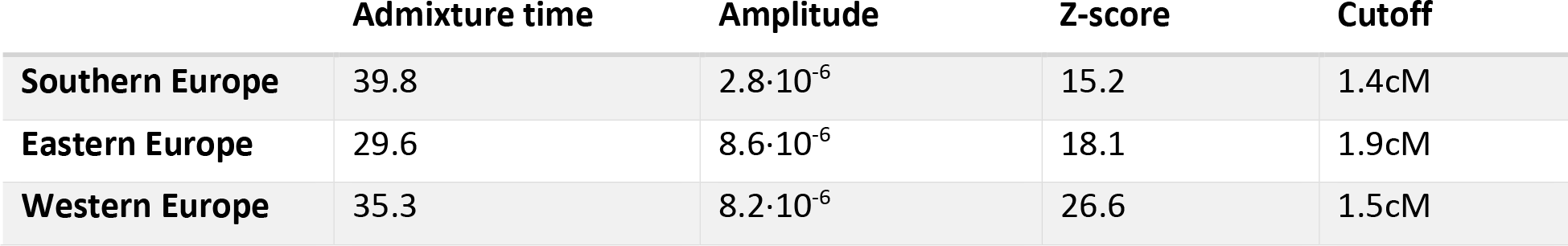
The results of inferring the AJ admixture time and sources using *Alder*. Admixture times are in generations. The parameters were inferred, for each European region, using *Alder’*s self-determined minimal distance cutoff (rightmost column), above which the admixture LD decay is fitted.

For a simple admixture history, the LD curve amplitude increases as the reference population becomes closer to the true ancestral source. The *Alder* results (Table 2) would thus suggest that Eastern and Western Europeans are closer to the true source of European gene flow into AJ, in contrast to the LAI-based results. However, when we ran *Alder* on simulated genomes with an admixture event, 30 generations ago, between Levant and Southern/Eastern/Western EU with respective ancestry proportions 50:35:12:3(%), the amplitudes were nearly identical to those of the real data, with the admixture times maintaining the same relative order and slightly overestimated at 34-41 generations ago. In fact, even simulations of pure Levant/Southern EU admixture resulted in roughly equal Southern vs. Eastern EU amplitudes. We thus conclude that, perhaps due to the complex admixture history in Southern Europe, *Alder* has no power to infer the true ancestral sources, whereas a model of predominantly Southern European contribution is still consistent with the data.

A similar situation was observed when inferring the ancestral tree topology using the *f_4_* statistics [48, 50] (Figures S4 and S5) as well as *TreeMix* [50] (Figure S6), both of which rely on the covariance of allele frequency between populations. We measured the *f_4_* statistic for the configuration (X,YRI;AJ,ME), where we used Yoruba (YRI) as an outgroup, and substituted different European regions for X (Figure S4A). The European region that gave the highest value of *f_4_*, Eastern Europe (closely matched by Western Europe), is theoretically the one closest to the true source of European gene flow. However, simulations with a dominant (or even exclusive) Southern European source resulted in highest *f_4_* values for Eastern Europe as well. [This discrepancy might be explained, at least partly, by a strong Middle-East to Southern EU migration event [51] (Figure S5)]. Therefore, those results are still consistent with a dominant Southern EU source for AJ. We also used the *f_4_* statistics to infer the fraction of European ancestry in AJ, as explained in Patterson et al. [48]. Assuming that the true source is Southern Europe, the EU ancestry proportion is given by *f_4_*(West-EU,YRI;AJ,ME)/*f_4_*(West-EU,YRI;South-EU,ME)≈67% (Figure S4B). However, when simulating 50% European ancestry, the *f_4_*-inferred fraction came out as 63%; thus, the inferred European ancestry proportion of 67% is consistent with the *RFMix*-based estimate of ≈53%.

We next ran *TreeMix* on AJ, Middle-East, the four European regions (West/East/South/Iberia), and YRI as an outgroup. The uncalibrated inferred tree (Figure S6) suggests that AJ split first, followed by Middle-Easterners and Europeans. *TreeMix* then predicted replacement of ≈42% of the Southern EU ancestry by Middle-Eastern migration, and ≈17% of the AJ ancestry by Eastern European migration, with the only other significant migration events coming from YRI and having lower magnitude. However again, simulations with a predominantly Southern European ancestry yielded nearly identical results (Figure S6). Interestingly, in simulations, *TreeMix* correctly estimated ≈13-14% Eastern EU ancestry in AJ when the true value was 12%, and almost no Eastern EU ancestry (≈2%) when none was simulated alongside Southern EU and ME ancestry; however, Eastern EU ancestry was erroneously estimated when the true simulated ancestry alongside Southern EU and ME was Western EU (16%).

In summary, we demonstrated that the raw results returned by *Alder*, thef-statistics, and *TreeMix* must only be interpreted in light of simulations. Using such simulations, the results were overall consistent with our previous model of an admixture event ≈35 generations ago involving predominantly Southern Europe, with minor contributions of either Western or Eastern Europe.

#### GLOBETROTTER analysis

Finally, we considered *GLOBETROTTER* [22], which can infer both the contribution of each ancestral source and the admixture time. The first step in *GLOBETROTTER* analysis is performed by *CHROMOPAINTER* [21], which determines the proportion of ancestry of each individual that is “copied” from each other individual in the dataset. Then, an *ancestry profile* for each population is reconstructed, representing the coefficients of a linear mixture of the copying vectors of each population [22, 23]. The inferred ancestry profile for AJ was 5% Western EU, 10% Eastern EU, 30% Levant, and 55% Southern EU. The combined Western and Eastern EU component is in line with our other estimates, as well as the dominance of the Southern EU component. However, the overall European ancestry, ≈70% (or ≈67% after calibration by simulations; Supplementary Text S3), is about 15% higher than the LAI-based estimate, as well as our previous results based on whole-genome sequencing [9]. Our detailed calibration simulations (Supplementary Text S3) demonstrate that evidence exists to support either estimate, suggesting that the true fraction of EU ancestry is midway, around ≈60% (see *Discussion*).

*GLOBETROTTER* is also able to directly infer admixture time and proportions, using the ancestry profiles, by assuming that the source groups could themselves be mixtures of the populations in the sample. A single admixture event was inferred for AJ (Supplementary Text S3), where the first source, comprising 36% of the total ancestry, was 46% Western EU and 53% Eastern EU. The second source (64% of the total ancestry) was 35% Southern EU and 65% Levant, and the inferred admixture date was 34 generations ago. Our simulations (Supplementary Text S3) show that the inferred total ≈22% of Southern EU ancestry is likely significantly underestimated (by ≈20 %-points), the overall inferred EU ancestry (here ≈58%) is accurate, and the inferred time is likely overestimated by ≈10 generations. With these adjustments, the results are broadly consistent with our conclusions so far. It remains open to explain the discrepancy between the results of the two modes of the program.

### Bounding possible historical models

We have so far provided multiple estimates for the ancestry proportions from each European source and the time of admixture events. We now attempt to consolidate those estimates into a single model and provide bounds on the model’s parameters. The results of all analyses (once calibrated by simulations) pointed to Southern Europe as the predominant source of European gene flow. At the same time, minor contributions from Western and/or Eastern Europe were also detected, with some analyses (IBD within AJ and between AJ and other sources, and *Globetrotter*) showing stronger support for an Eastern European source. Based on historical plausibility, these admixture events must have necessarily happened at different times, implying multiple historical events. The inferred admixture time, when modeled by a single event, was between ≈24-40 generations ago by the methods we examined (calibrated mean segment length and ancestry proportions, *Alder*, and *Globetrotter*), very close to the time of the AJ bottleneck, previously estimated to ≈25-35 generations ago [9, 16]. Therefore, admixture must have occurred both before and after the bottleneck, with the IBD and *Alder* analyses suggesting that the Eastern European admixture was more recent.

Based on these arguments, we propose that a minimal model for the AJ admixture history includes substantial pre-bottleneck admixture with Southern Europeans, followed by post-bottleneck admixture on a smaller scale with Western or, more likely, Eastern Europeans. The estimates for the total European ancestry in AJ range from ≈49% using our previous whole-genome sequencing analysis [9], to ≈53% using the LAI analysis here, and ≈67% using the calibrated *Globetrotter* analysis. The proportion of Western/Eastern European ancestry was estimated between ≈15% (*Globetrotter* and the LAI-based localization method), and, if identified as the source of the post-bottleneck admixture, 23% (the IBD analysis). Therefore, the proportion of the Southern European (presumably pre-bottleneck) ancestry in AJ is between ≈26% to ≈52% (corresponding to [33,61]% ancestry at the time of admixture). Given those bounds, along with the admixture time estimate based on a single event (24-40 generations ago), we could derive an equation to constrain the admixture times of the pre- and post-bottleneck events (*Methods*). We assumed that the post-bottleneck admixture event happened 10-20 generations ago; for the upper bound, we allowed ≈10 generations since the bottleneck for the effective population size to reach thousands, at which point barely any within-AJ IBD segments descend from these migrants (see the IBD analysis above and *Methods*); for the lower bound, no mass admixture events are known in the past 2-3 centuries of AJ history [52]. The results (*Figure 6*) show that given these constraints, the pre-bottleneck admixture time is between 24-53 generations ago. Our proposed model is shown in Figure 7.

**Figure 6.**
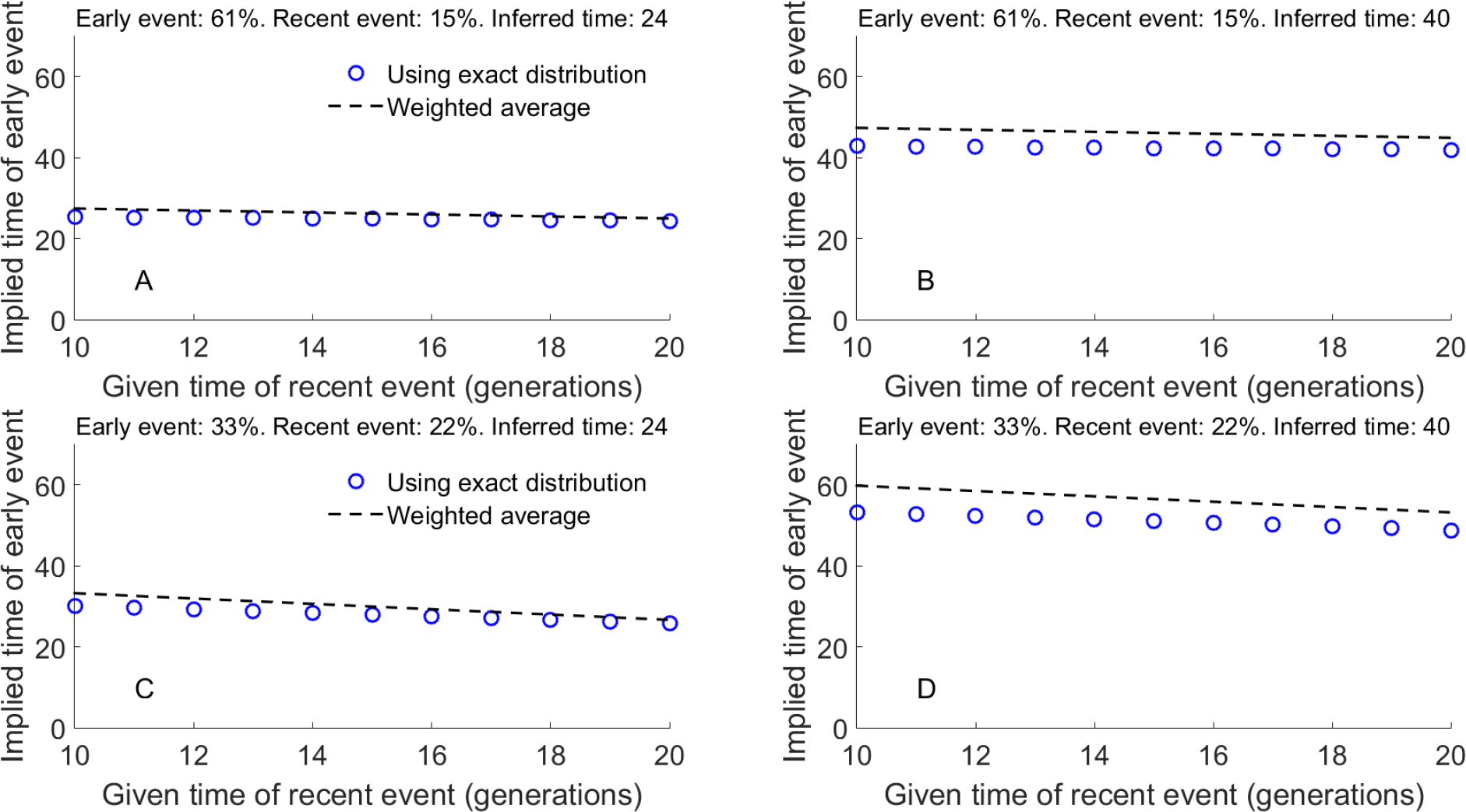
The relationship between two admixture times, given bounds on other admixture parameters. In this model, two populations (**A** and **B**) mixed at time *t*_1_ (*early event*; the proportion of ancestry contributed by population **A**, *q*, is indicated in the title of each panel). At a more recent time, *t*_2_ (*recent event*), migrants from *A* replaced another proportion *μ* of the admixed population (proportions also indicated in the titles). In each panel, we assumed that *q* and *μ* are known, as is the admixture time inferred using a pulse admixture model (titles). Under these assumptions, and using Equation (6) in *Methods*, we plotted the time of the early event (*t*_1_) vs the time of the recent event (*t*_2_; blue circles). The weighted average (dashed lines) is a simple approximation, in which the time inferred under the pulse model is a weighted average of *t*_1_ and *t*_2_, weighted by the admixture proportions *q* and *μ*, respectively. In the context of the Ashkenazi Jewish admixture history, population **A** is European and **B** is Middle-Eastern. Panels (A)-(D) represent the bounds on (*i*) the admixture time inferred under a pulse model (24-40 generations ago); (*ii*) the admixture proportions at the early and recent events (33-61% and 15-23%, respectively), as described in the main text; and (*iii*) the time of the recent admixture event (10-20 generations). The results show that (*i*) the weighted average is a reasonable approximation, though the pulse admixture time is influenced more by the early event, perhaps as it results in more admixture tracts; and (*ii*) the most extreme values of the early AJ event are 24 and 53 generations ago. The lower bound correspond to the lowest value of the inferred (single event) admixture time, the highest value of the time of the recent admixture event, and the largest contribution of the early event to the overall admixture proportions (and vice versa for the upper bound).

**Figure 7.**
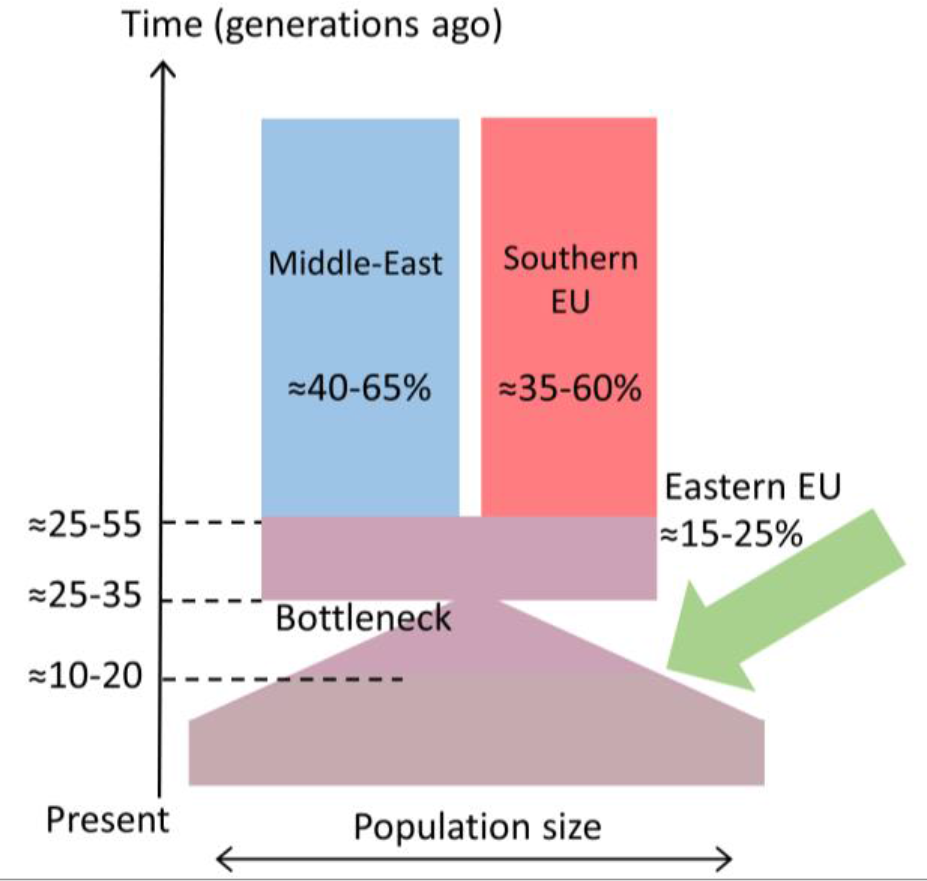
A proposed model for the recent AJ history. The proposed intervals for the dates and admixture proportions are based on the spectrum of estimates obtained by the different analyses, as described in the text.

## Discussion

### Summary and lessons

The ethnic origins of Ashkenazi Jews have fascinated researchers for over a century [53, 54]. The availability of dense genotypes for thousands of AJ individuals, along with the development of sophisticated analysis tools, established close relations between AJ and other Jewish groups, as well as putative European and Middle-Eastern origins [4–8, 25]. Here we attempted, for the first time, to create a detailed portrait of the admixture events experienced by AJ during their dwelling in Europe. To this end, we used previously generated genome-wide array data for AJ, European, and Middle-Eastern populations (*Table 1*), as well as a variety of methods from the population genetics toolbox, including some newly developed techniques.

Before discussing the historical implications of our results, we point out two general lessons that emerge from the analysis. The first lesson is that AJ genetics defies simple demographic theories. Hypotheses such as wholly Khazar or wholly Middle-Eastern origins have already been disqualified [4–7, 18], but even a model of a single Middle-Eastern and European admixture event cannot account for all of our observations, and the actual admixture history might have been highly complex. Moreover, due to the genetic similarity and complex history of the European populations involved (particularly in Southern Europe [51]), the multiple paths of AJ migration across Europe [10], and the strong genetic drift experienced by AJ in the late Middle Ages [9, 16], there seems to be a limit on the resolution to which the AJ admixture history can be reconstructed.

The second lesson is the importance of evaluating the results of off-the-shelf tools using simulations of the specific populations under study. When simulating Middle-Eastern and European admixture (particularly Southern European), we found many tools to be of limited utility (see, e.g., the section on *Alder*, *f*-statistics, and *TreeMix*, the *Methods* section on *ADMIXTURE*, and Supplementary Texts S1 and S2 on *LAMP* and *PCAMask*). Further, while we were eventually able to extract useful information off *RFMix’*s local ancestries, the raw results were not very accurate: the accuracy per SNP was only ≈70-80%, the mean segment length was more than twice than expected, and the variance of the ancestry proportion per chromosome was underestimated. When jointly analyzing LAI and IBD sharing, the inferred proportion of IBD segments that were either false or had random LAI was as high as ≈35% ((1-λ) in *Methods*), although fortunately, we were able to account for that in the model.

### Historical model and interpretation

Our model of the AJ admixture history is presented in *Figure 7*. Under our model, admixture in Europe first happened in Southern Europe, and was followed by a founder event and a minor admixture event (likely) in Eastern Europe. Admixture in Southern Europe possibly occurred in Italy, given the continued presence of Jews there and the proposed Italian source of the early Rhineland Ashkenazi communities [3]. What is perhaps surprising is the timing of the Southern European admixture to ≈31-52 generations ago, since Jews are known to have resided in Italy already since antiquity. This result would then imply no gene flow between Jews and local Italian populations until the turn of the millennium, either due to endogamy, or because the group that eventually gave rise to contemporary Ashkenazi Jews did not reside in Southern Europe until that time. More detailed or alternative interpretations are left for future studies.

Recent admixture in Northern Europe (Western or Eastern) is consistent with the presence of Ashkenazi Jews in the Rhineland since the 10^th^ century and in Poland since the 13^th^ century. Evidence from the IBD analysis suggests that Eastern European admixture is more likely; however, the support is not decisive. An open question in AJ history is the source of migration to Poland in late Medieval times; various speculations have been proposed, including Western and Central Europe [2, 10]. The uncertainty on whether gene flow from Western Europeans did or did not occur leaves this question open.

### Caveats and future work

The historical model we proposed is based on careful weighting of various methods and simulations, and we attempted to account for known confounders. However, it is possible that some remain. One major concern is the effect of the narrow AJ bottleneck (effective size ≈300 around 30 generations ago [9, 16]) on local ancestry inference and other methods, in particular *TreeMix* and *f*-statistics, but also more complex models such as *GLOBETROTTER*, for which the effect of drift is less obvious. Another general concern is that while we assumed the different methods provide independent pieces of evidence, they might have modeled the same features of the data, or worse, the same artifacts. We generally attempted to avoid the effect of the bottleneck (*Methods*) as well as select methods orthogonal to each other, but some issues may have remained.

Another caveat is that our estimation of the two-wave admixture model is based on heuristic arguments (the multiple European sources and the differential ancestry at IBD segments), and similarly for the admixture dates. The IBD analysis itself relies on a number of assumptions, most importantly that the error in LAI is random and distributed according to the total ancestry fractions. Those fractions are themselves difficult to estimate, as can be seen by the discordance between *RFMix* and *GLOBETROTTER* (53% and 70% EU fraction, respectively).

While our AJ sample is extensive, our reference panels, assembled from publicly available datasets, are incomplete. Specifically, sampling is relatively sparse in North-Western and Central Europe, and sample sizes in Eastern Europe are rather small (10-20 individuals per population). Our partitioning of the sample to broad geographic regions is somewhat arbitrary, grouping together heterogeneous populations. In addition, we did not consider samples from the Caucasus or from non-Ashkenazi Jewish communities, and this could have slightly affected the analysis (although likely not significantly [5]). Similarly, we neglected any sub-Saharan ancestry, even though Southern European and Middle-Eastern populations (including Jews) are known to harbor low levels (≈5-10%) of such ancestry from earlier migrations [49, 55]. Finally, a commonly overlooked problem is that a reference population currently representing one geographic region might have migrated there recently, thereby misrepresenting the true geographic origins of the ancestral sources of the admixed population studied. However, this is not expected to be a major concern here, where our geographic regions span very large areas.

The admixture history of Ashkenazi Jews thus remains a challenging and partly open question. To make further progress, the natural next step is to use sequencing data. Whole-genomes are now available for several European populations (e.g., [56]) as well as for Ashkenazi Jews [9] and some Middle-Eastern groups [57]. Our results demonstrate that the accuracy of LAI is expected to increase for sequencing data (not shown), and similar conclusions were made for other analysis tools (e.g., [58]). Additionally, the availability of whole-genomes will make possible analyses based on the allele frequency spectrum in AJ and other populations. At the same time, denser sampling of relevant European and Middle-Eastern populations (mostly from Central and Eastern Europe) will be required in order to refine the geographic source(s) of gene flow.

Beyond data acquisition, we identify three major methodological avenues for future research into AJ admixture. First, any improvement in the accuracy of local ancestry inference will translate into improved power to resolve admixture events, in particular for events within Europe. Second, since AJ admixture history was complex, new methods will have to be developed and applied for the inference of continuous and multi-wave admixture histories (e.g., [35, 59]). At the same time, inference limits will have to be established for events temporally or geographically near, as we began to develop here (Supplementary Text S4; see also [40]). Finally, one may use the signal in the lengths of IBD segments shared between AJ and other populations to construct an admixture model (e.g., as in [60]), which may be less prone to noise than the LAI-based estimate, provided that we can reliably detect shorter segments than currently possible.

## Methods

### Data collection

After merging genotypes from all sources (*Table 1*, lifting over to hg19 whenever necessary), cryptic relatives were removed by first detecting IBD segments (*Germline* [45]) and then removing one of each pair of individuals sharing more than 300cM. Individuals with a non-Ashkenazi genetic ancestry (defined to share less than 15cM, on average, with other AJ) were also removed. Other standard QC measures (carried out in *Plink* [61]) included removal of SNPs or individuals with high no-call rate and eliminating SNPs with an ambiguous strand assignment. The genotypes were phased using *Shapeit* [62].

### Local ancestry inference using *RFMix*

*RFMix* was run using the TrioPhased option (see Supplementary Text S1) and the generation parameter set to 30. Other parameters were set to default values. In each analysis involving the AJ individuals, we used a random subset of 400 or 500 individuals (out of overall 2540) in order to save computational time. We did not use the expectation maximization (EM) option, as simulations of ME/EU admixture demonstrated that inference accuracy was not improved by running the highly time-consuming EM step. Additionally, the EM step makes iterative use of the admixed (Ashkenazi) genomes themselves in order to supplement the reference panels, thereby potentially introducing biases due to the excessive haplotype sharing in AJ. We therefore decided not to use EM for our subsequent analysis.

#### Balancing the reference panels

To minimize biases in local ancestry inference, we ensured an equal number of European and Middle-Eastern individuals in the reference panel, as well as an equal number (30) of individuals from each subcontinental European region (South, West, East, and Iberia). We then used the same reference panel both for testing our simulations and for the AJ data, but the reference panel did not include the genomes used to create the simulated individuals (20 from each EU region and 20 from the Levant region). An initial inspection of our geographic localization pipeline demonstrated that Iberia had a much lower likelihood compared to the other regions. We thus removed Iberia from our reference panel for inference, which allowed us to significantly increase the number of individuals used in the remaining regions (as Iberia had the smallest number of available genomes). Our final reference panel consisted of 273 EU and 273 ME individuals: 91 Eastern European, 91 Western European, 91 Southern European, 70 Druze, 77 Southern Middle-Eastern, and 126 Levantine individuals.

#### Global ancestry proportions

To infer the global ancestry proportions from *RFMix*, we used the proportion of SNPs classified as European/Middle-Eastern. Global ancestry estimates were also inferred using *ADMIXTURE* [41] (default parameters), either supervised or unsupervised. Surprisingly, when we tested simulated Southern European/Levantine admixed genomes, the unsupervised mode yielded more accurate ancestry proportions than the supervised mode. We also found that inferring global ancestry using *RFMix* outperformed *ADMIXTURE*.

### Simulation details

For each admixed individual, we assumed that admixture (from all source populations) occurred at a single generation. The admixture parameters are the ancestry proportion contributed by each source and the admixture time *G* (generations ago). We generated a haploid chromosome sequentially until reaching the end of the chromosome. The ancestry of each chromosomal segment was randomly selected, using the weight of each source (i.e., its ancestry proportions). We then randomly selected a chromosome from the chosen source population, and drew the segment length (in cM) from an exponential distribution with rate *G*/100. A haploid set of 22 chromosomes was then created for each individual. Diploid individuals were constructed by randomly pairing two sets of haploid chromosomes.

### IBD sharing analysis

#### IBD segment detection

Five hundred random AJ individuals were selected for the IBD analysis. IBD segments were detected using *Germline* [45] with parameters *bits*=64, *err*_*hom*=1, *err*_*het*=1, and a minimum length of 3cM. The detected segments were filtered with *Haploscore* [46] (cutoff 2) as well as eliminating segments with more than 5% overlap with sequence gaps. In the analysis of the ancestry of the segments, we eliminated 0.25cM at each end of each segment to account for misidentification of their boundaries.

#### Ancestry of IBD segments

Denote by *p*_EU_ the genome-wide proportion of European ancestry in the AJ genomes, and assume it is known (e.g., ≈53%, as obtained from the LAI (*RFMix*) analysis). The goal of the IBD analysis is to compare *p*_EU_ to the proportion of EU ancestry in the IBD segments. Complicating the analysis are (*i*) that the reported IBD segments are between diploid genomes (even though sharing is between single haplotypes); and (*ii*) errors in IBD detection and local ancestry inference. Nevertheless, the genome-wide expected effect of these confounders could be completely accounted for. To see this, denote by λ the proportion of IBD segments that are both real and whose inferred local ancestry is correct. The remaining segments (proportion 1-λ) are either not IBD or their inferred local ancestry is random. In both cases, the local ancestry assignment is EU with probability *p*_EU_ and ME with probability 1-*p*_EU_. Next, define the observed IBD ancestry matrix **A**_obs_, whose rows correspond to the ancestry of the segment at individual 1 (with three possibilities: hom-EU, het, and hom-ME) and whose columns correspond to individual 2. Each entry in the matrix corresponds to the proportion of genetic material in IBD segments (genome-wide, in cM) where the two individuals have the given ancestries. The matrix **A**_rand_ is similarly defined, for either random regions or random local ancestry assignment. **A**_rand_ has expectation

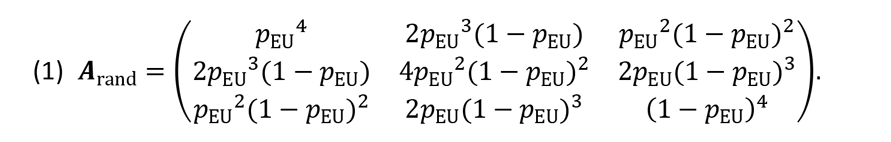

(To simplify notation, and since there is no ambiguity, we use the same symbol for the matrix and its expectation.) For true IBD regions, we assume that all IBD segments descend from a common ancestor that lived around the time of the bottleneck (see below for justification). We denote the genome-wide EU ancestry during the bottleneck as *f*_EU_, which could be different than *P*_EU_: for example, a wave of post-bottleneck European gene flow would imply *f*_EU_ < *p*_EU_. At IBD segments, the two individuals have, by definition, only three independent chromosomes (the one shared, and the other chromosome at each individual). The shared chromosome will be European with probability *f*_EU_, while the two other chromosomes will be European with probability *p*_EU_. Denote by **A**_IBD_ the ancestry matrix at IBD segments, and write its expectation as

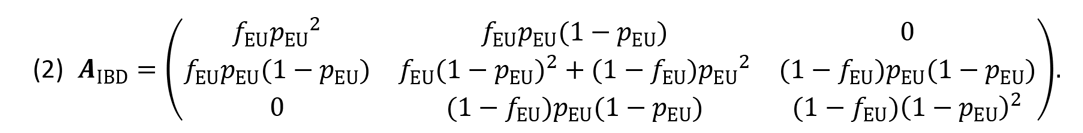

Note that no IBD segments have hom-EU ancestry for one individual and hom-ME ancestry for the other. Finally, we have

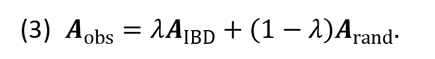

The noise level λ is then estimated by matching the (hom-EU/hom-ME) elements in **A**_obs_ and **A**_rand_, since none of these elements depends on the unknown *f*_EU_. Given λ, the empirical AIBD can be computed from Eq. (3). We then estimate *f*_EU_ by minimizing the sum of absolute differences between the empirical and theoretical elements of **A**_IBD_. Note that the calculation above relies on the assumption that the ancestry of segments with false positive IBD or uninformed LAI is random (with EU ancestry proportion *p*_EU_). Another assumption is that given that a site is in an IBD segment, it coalesces around the time of the bottleneck. The exact posterior distribution of the coalescence time is given by (e.g., [16, 63])

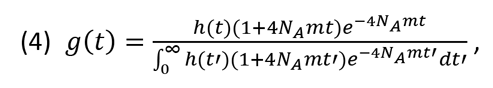

where *m* is the minimal segment length, *N_A_* is the ancestral population size, *h*(*t*) is the coalescence probability per generation (or the inverse population size when scaled by *2N_A_*), and the time *t* is scaled by *2N_A_*. For a bottleneck of ≈300 individuals around 30 generations ago followed by rapid expansion as inferred for AJ [9, 16], we find that coalescence times are narrowly distributed, with ≈90% of events taking place within 15 generations of the bottleneck. This suggests that the ancestry of IBD segments reflects predominantly the ancestry during the generations close to the bottleneck. [Information on deviations from this assumption is encoded in the lengths of the segments and may be modeled in future work.]

To determine the pre- and post-bottleneck admixture proportions, we assume a model of pre-bottleneck admixture with proportions *f*_EU_:(1−*f*_EU_) and a post-bottleneck wave of European gene flow of magnitude *μ_EU_*. The total proportion of EU ancestry, *p*_EU_, can be written as *p*_EU_=*μ*_EU_ + (1 − *μ*_EU_) · *f*_EU_. Given the observed *p*_EU_ and the estimated *f*_EU_, *μ*_EU_ can be obtained.

#### *Alder*, *f*-statistics, and *TreeMix* analyses

We ran *Alder* [47] with default parameters (including automatic detection of the minimal length cutoff), and with two reference populations. *f_4_* statistics were computed using the implementation in the *TreeMix* package [50]. The *TreeMix* analysis itself was run with default parameters, except a block size (− k) of 500 (corresponding to ≈5MB, beyond the extent of typical LD).

#### *GLOBETROTTER* analysis

On both simulations and real AJ data, *GLOBETROTTER* was run with default settings, as given in the example distributed with the program. For completeness, when generating only ancestry profiles (the proportion of ancestry contributed by each reference population), the key parameters were set to prop.ind=1 and num.mixing.iterations=0. When inferring both admixture events and proportions, we used boostrap.num=20, props.cutoff=0.001, and num.mixing.iterations=5. To save computational time when running *GLOBETROTTER* on the real data, we used a random subset of 200 AJ individuals.

### Inferring admixture times using the distribution of ancestry proportions

Several methods have been recently proposed for the estimation of historical admixture times. Johnson et al. [19] fitted the number of ancestry switches; Pugach et al. [64] matched simulations to the typical segment length, as estimated from a wavelet transform of the local ancestry along the genome; and Pool and Nielsen [65], as well as Gravel [35], fitted the distribution of segment lengths. However, these methods require an accurate identification of the boundaries of admixture segments, which is not always easy, in particular for computationally phased data. Reich and colleagues [47–49] fitted the decay of admixture linkage disequilibrium (LD) with genetic distance (see main text), but their method can be confounded by background LD. Hellenthal et al. [22] recently proposed a promising approach based on the probability of two fixed loci to have given ancestries. Admixture parameters can also be inferred using more general demographic inference methods, e.g., based on the allele frequency spectrum [66, 67] or segment sharing [60]; however, to use these methods one must specify and infer a model for the entire history. Recently, Rosenberg and colleagues [39, 68], Liang and Nielsen [69], and Gravel [35], derived analytical results for the moments of the ancestry proportion, namely the portion of the chromosome that descend from a given ancestry. These ancestry proportions can be reliably inferred (e.g., [41, 70]), and the derived moments have been used for admixture time inference (e.g., [36, 37]). However, these methods do not make use of the information available in the entire distribution. We therefore sought to derive this distribution.

Our method assumes a simple admixture model, where the admixed population under investigation formed *t* generations ago as a result of merging of populations **A** and **B**, and where the proportion of ancestry contributed by **A** and **B** was *q* and 1 − *q*, respectively. Each locus in a chromosome of a present-day admixed individual can trace its origin to **A** or **B** with probabilities *q* and 1 — *q*, respectively. We assume that lineages break apart along the chromosome due to recombination, at rate *t* per Morgan. Ignoring genetic drift and constraints imposed by the underlying shared pedigree [71], we assume that following recombination, the new source population is selected at random. Therefore, a recombination event will lead to a change of ancestry from *A* to *B* with probability 1 − *q* and from **B** to **A** with probability *q*. The lengths of the chromosomal segments with **A** ancestry will therefore be exponentially distributed with rate (1 − *q*)*t*, and similarly for the **B** segments (rate *qt*) [35]. We neglect the first generation after admixture where **A** and **B** segments do not yet mix [35]. As pointed out by Liang and Nielsen [40], the assumption of independent and exponentially distributed segment lengths breaks down for very short and very long times, due to the effect of the underlying pedigree and the accumulation of genetic drift, respectively. However, for admixture in a population such as Ashkenazi Jews (admixture time around 10-80 generations, and population size much larger than the number of generations even at the bottleneck), segment lengths should be very well approximated by independent exponentials.

Given a chromosome of length *L* (Morgans), the ancestry along the chromosome can be modeled as a two-state process with states **A** and **B**, and with the distribution of segment lengths in each state given above. We are interested in the distribution of *x*, the fraction of the chromosome in state **A**. Adopting a result of Stam [38], the desired distribution is given by

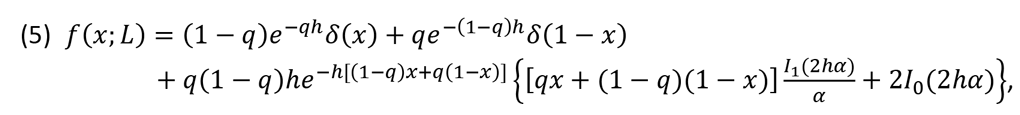

where 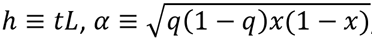 and *I*_0_ and *I*_1_ are the modified Bessel functions of the first kind of order 0 and 1, respectively. Note the delta functions at *x* = 0 and *x* = 1, corresponding to the probability of the entire chromosome to have **B** only or **A** only ancestry, respectively. The mean ancestry proportion satisfies 〈*x*〉 = *m*, as expected. The variance is given by

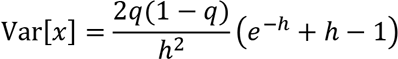

in agreement with Eq. (A16) in [35].

In practice, in the absence of trios or pedigree information, phase switch errors are abundant and hence it is difficult to accurately determine the ancestry proportion per chromosome. However, it is still possible to determine the diploid ancestry proportion, *y* = (*x*_1_+*x*_2_)/2. Given that homologous chromosomes have independent histories, its distribution, *f_d_*(*y;L*), can be computed from Eq. (5) by convolution. Suppose we are now given the diploid ancestry proportions *y_ij_* for individuals *i* = 1, …,*n* and for chromosomes *j* = 1,…,22 (where each chromosome has length *L_j_*). Assuming that chromosomes are independent both within and between individuals, the likelihood of the data is given by

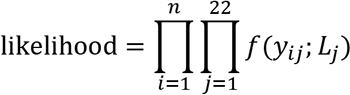

We can then maximize the likelihood using a simple grid search over *q* and *t*. Simulation results with perfect knowledge of segments boundaries demonstrated that the method can infer correctly both *q* and *t* with very small variance (Figure S2), although the variance increases with *t*, as expected. Coalescent simulations followed by inference of ancestry proportions using *ADMIXTURE* [41] and application of our method demonstrated again high accuracy, at least as long as the **A** and **B** populations were sufficiently diverged (not shown). However, when **A** and **B** were closely related, the distributions of the true and inferred ancestry proportions differed; in this case, simulations must be used to calibrate the inferred time (see main text).

Applying the method to the AJ ancestry proportions, we obtain a point estimate of *q* = 0.55 and *t* = 22 generations. Resampling the ancestry proportions 25 times over individuals (for each chromosome separately), we obtained bootstrap estimates of *q* = 0.547 α 0.009 (mean α standard deviation; range 0.53 − 0.56) and *t* = 24.1 α 2.7 (range 20 − 31), although we note that the systematic error due to LAI errors is much higher than the sampling error.

We also considered a more complex historical model with an additional admixture event. Under this model, populations A and B had merged *t*_1_ generations ago, contributing proportions *q* and 1 − *q* to the admixed population. Then, *t*_2_ (< *t*_1_) generations ago, migrants from population **A** have replaced a proportion *μ* of the gene pool of the admixed population. No other events then take place until the present. Using the Markov process representation of the admixture process of Gravel [35], and using techniques of renewal theory, we were able to derive the distribution of the lengths of the A and B segments, which depend, in a complex way, on (*t*_1_, *t*_2_, *q*, *μ*). We then obtained an implicit expression for the distribution of the ancestry proportion over a chromosome. (More specifically, we obtained the Laplace transform of that distribution with respect to the chromosome length.) Mathematical details are given in the Supplementary Text S4. However, we observed that the distribution of ancestry proportions generated from the double admixture model often fits well to the pulse model (Supplementary Text S4), and therefore, we did not use our theoretical results for direct inference.

Nevertheless, these results are useful for understanding the range of double admixture models that will be inferred as identical pulse admixture events. Specifically, under double admixture, the distribution of B segments is exponential with rate *r* = *t*_1_ — (1 − *q*)(*t*_1_ − *μt*_2_), and the proportion of **B** ancestry is *M* = (1 − *q*)(1 − *μ*). Since for pulse admixture *T* generations ago, *r* = (1 − *M*)*T*, then the inferred time *T* under a pulse model satisfies

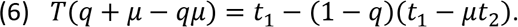

Given *T*, Eq. (6) imposes a constraint on the parameters of the model, in particular if *q* and *μ* can be independently estimated, as in our case.

## Acknowledgements

We thank Harry Ostrer for proposing the analysis of local ancestry in Ashkenazi Jews and Iain Mathieson and Shaul Stampfer for discussions. We thank financial support from the Hebrew University of Jerusalem and The Barouh and Channah Berkovits Foundation (SC).

## Supplementary Figures

**Figure S1.**
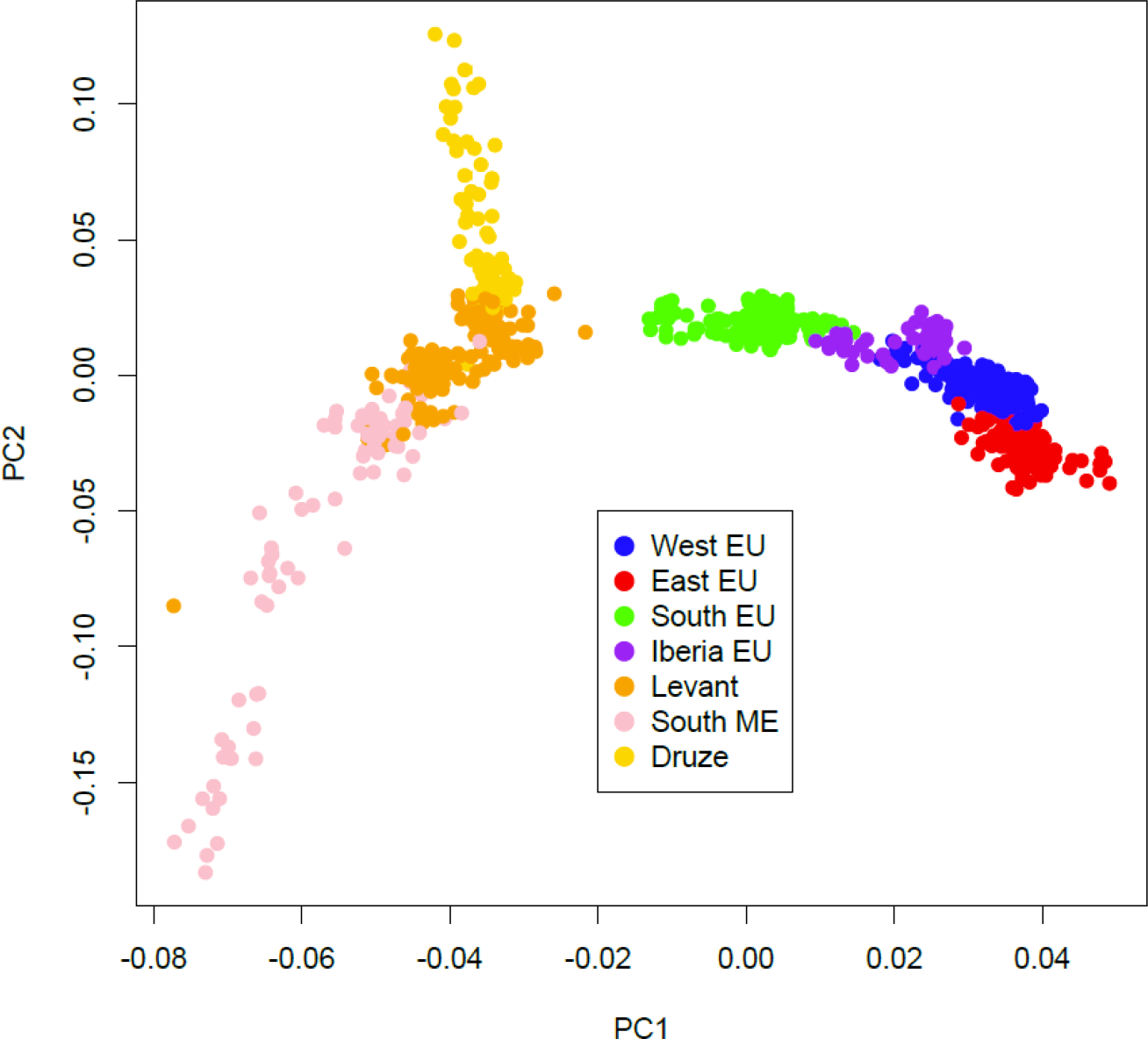
Principal Component Analysis (PCA) of the European and Middle-Eastern samples used as reference panels in our study. The analysis was performed by *SmartPCA* [72] using default parameters (except no outlier removal). The populations included within each region (as indicated in the legend) are listed in *Table 1* of the main text. The PCA plot supports the partitioning of the European and Middle-Eastern populations into the broad regional groups used as reference panels.

**Figure S2.**
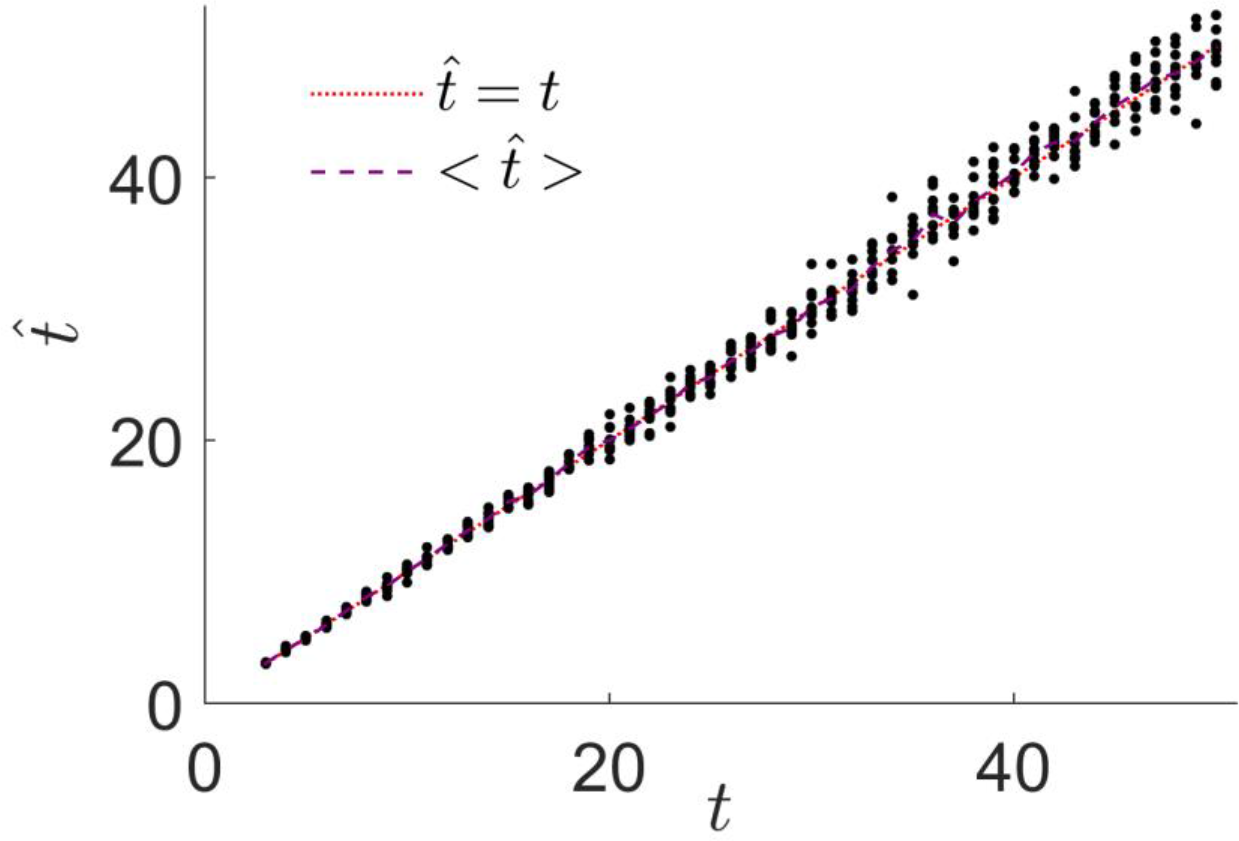
Inference of admixture time using the distribution of ancestry proportions. We simulated an admixture pulse history under the Markovian Wright-Fisher model of Gravel, *Genetics* **191**, 607 (2012). The model assumes that the 2*N* haploid chromosomes in the current generation are formed by following a Markovian path within the 2*N* chromosomes of the previous generation. Ancestry changes occur as a Poisson process with rate 1 (per Morgan). Each chromosome in the first generation is assigned to population **A** or **B** with probabilities *q* and 1 − *q*, respectively, and the evolution of the chromosomes is traced for *t* generations. We used *q* = 0.5, *L* = 2M, and *N* = 2500, and varied *t*. Ancestry proportions from pairs of chromosomes were averaged to generate diploid individuals. We then set the inferred *q* to the mean **A** ancestry, and used the distribution of ancestry proportions over the simulated individuals (see *Methods* in the main text) to infer the admixture time *t*. Each dot in the plot shows the inferred time, 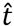 for one simulation. The dotted red line corresponds to 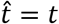 and the dashed purple line to the mean inferred time, 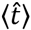.

**Figure S3.**
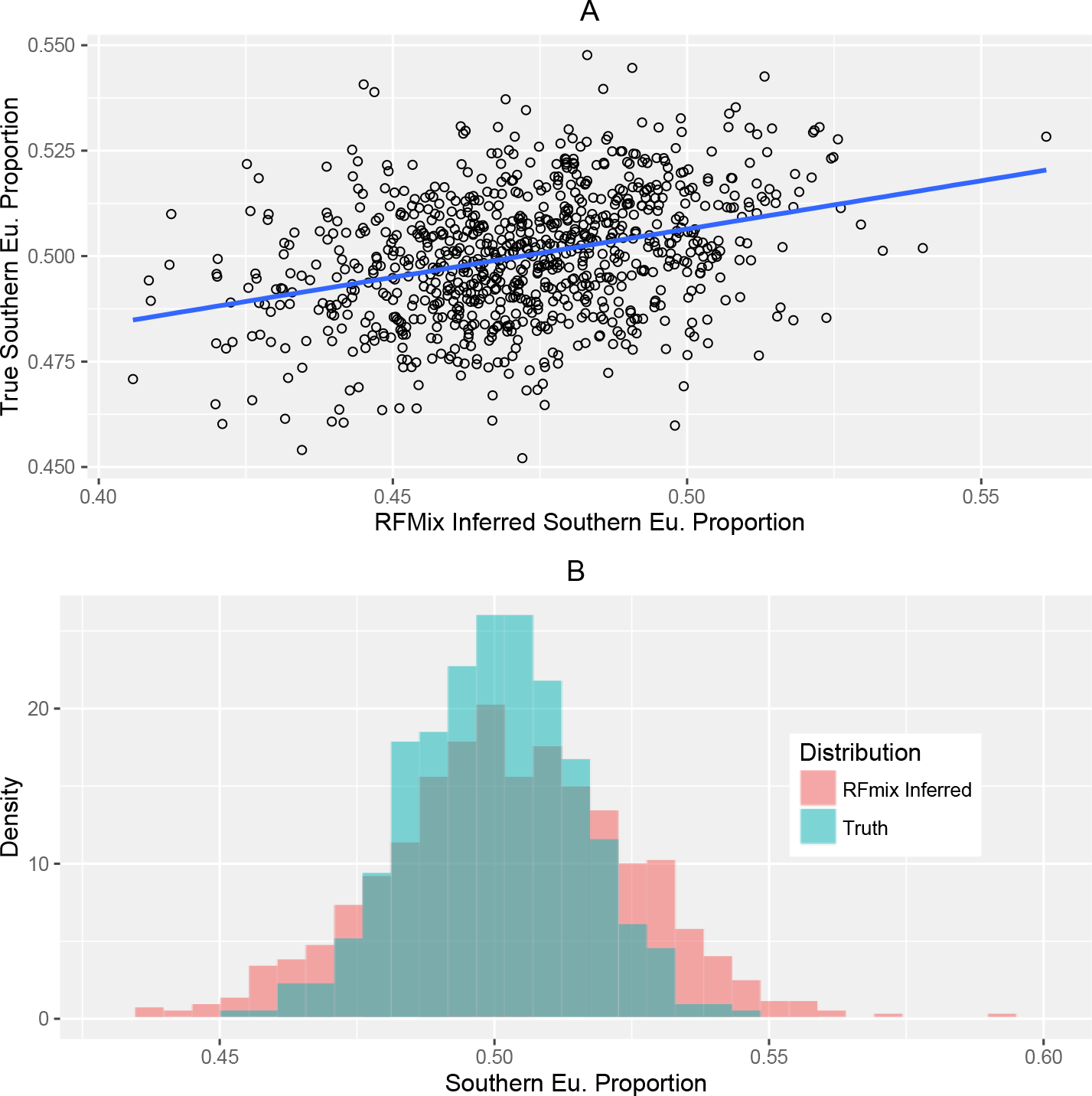
Simulation of 870 admixed individuals with 50% Southern European ancestry, 50% Levantine ancestry, and admixture time 30 generations ago. (A) Simulated vs *RFMix*-inferred Southern European ancestry proportion (*r*^2^ = 0.11). (B) The distributions of the simulated and *RFMix*-inferred ancestry proportions. The inferred proportions have larger variance than the true one, as well as a slightly lower mean (difference 0.03; for visualization, we shifted the *RFMix*-inferred distribution to match the true mean). A similar analysis with a European component being entirely Western European resulted in a much higher correlation (*r*^2^ = 0.5), although with a somewhat larger bias (0.11 above than the true mean).

**Figure S4.**
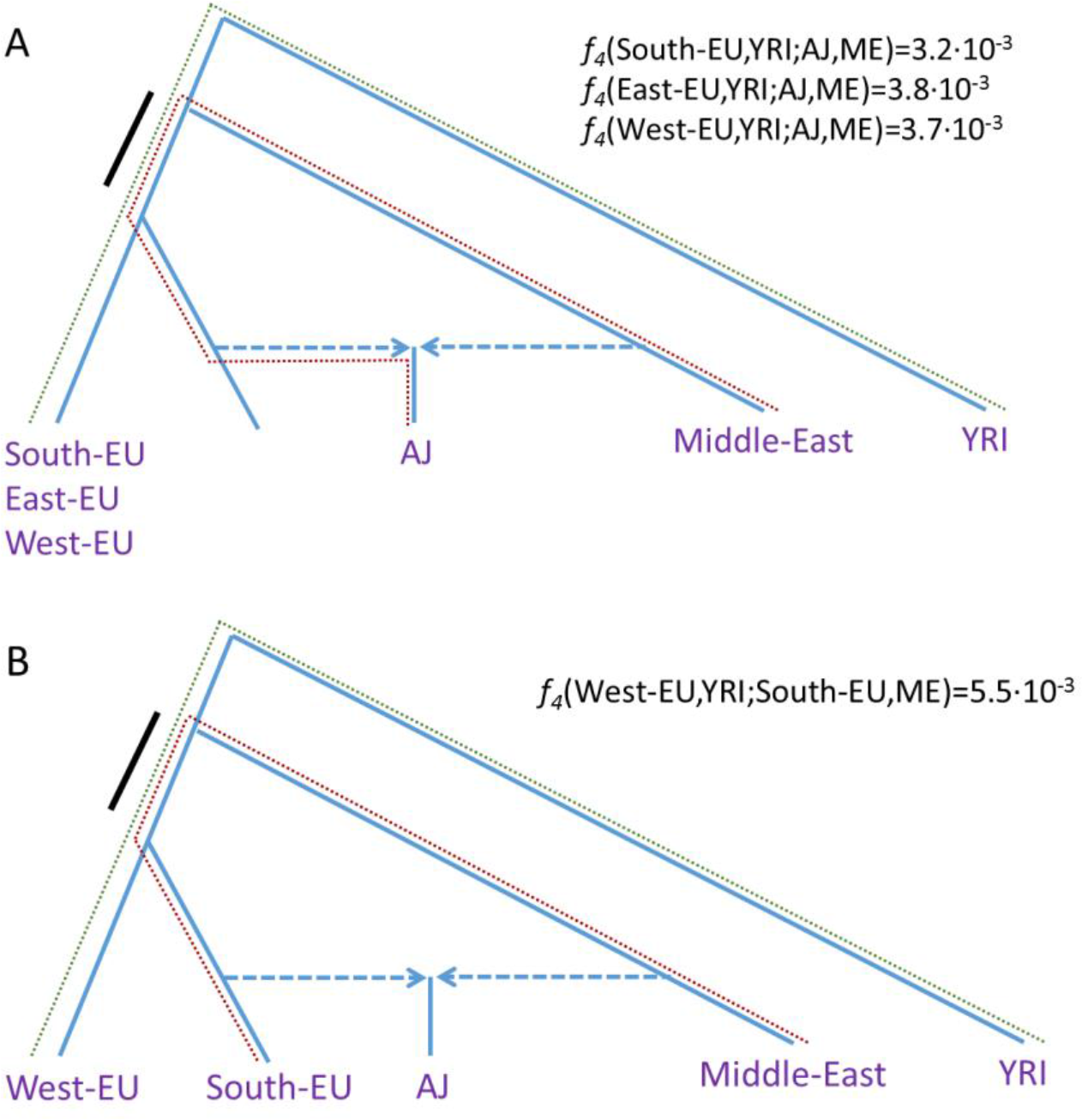
*f_4_* statistics and potential tree topologies for the AJ history. The method is based on Patterson et al. [48]. (A) Determining the likely source of European gene flow into AJ. The statistic *f_4_*(X,YRI;AJ,ME) compares the amount of shared ancestry (solid black bar) between the lineages connecting the European population X and Yoruba (green dashed lines) and the lineages connecting AJ and Middle-Easterners (red dashed lines). The closer population X is to the true source of gene flow, the larger should be the *f_4_* statistic. However, while we found higher values of *f_4_* for Western and Eastern Europe, simulations show that this pattern is reproduced even under simulations with a predominantly Southern European source. (B) Estimating the European ancestry fraction. This is similar to (A), except that we computed the statistic *f_4_*(West-EU,YRI;South-EU,ME) (assuming that Southern Europe is the true source of European gene flow). As explained in Patterson et al. [48] (Figure 2C therein), under the assumed tree topology, the ratio between the *f_4_* statistics in (A) (with X=West-EU) and (B) should equal the fraction of European ancestry in AJ.

**Figure S5.**
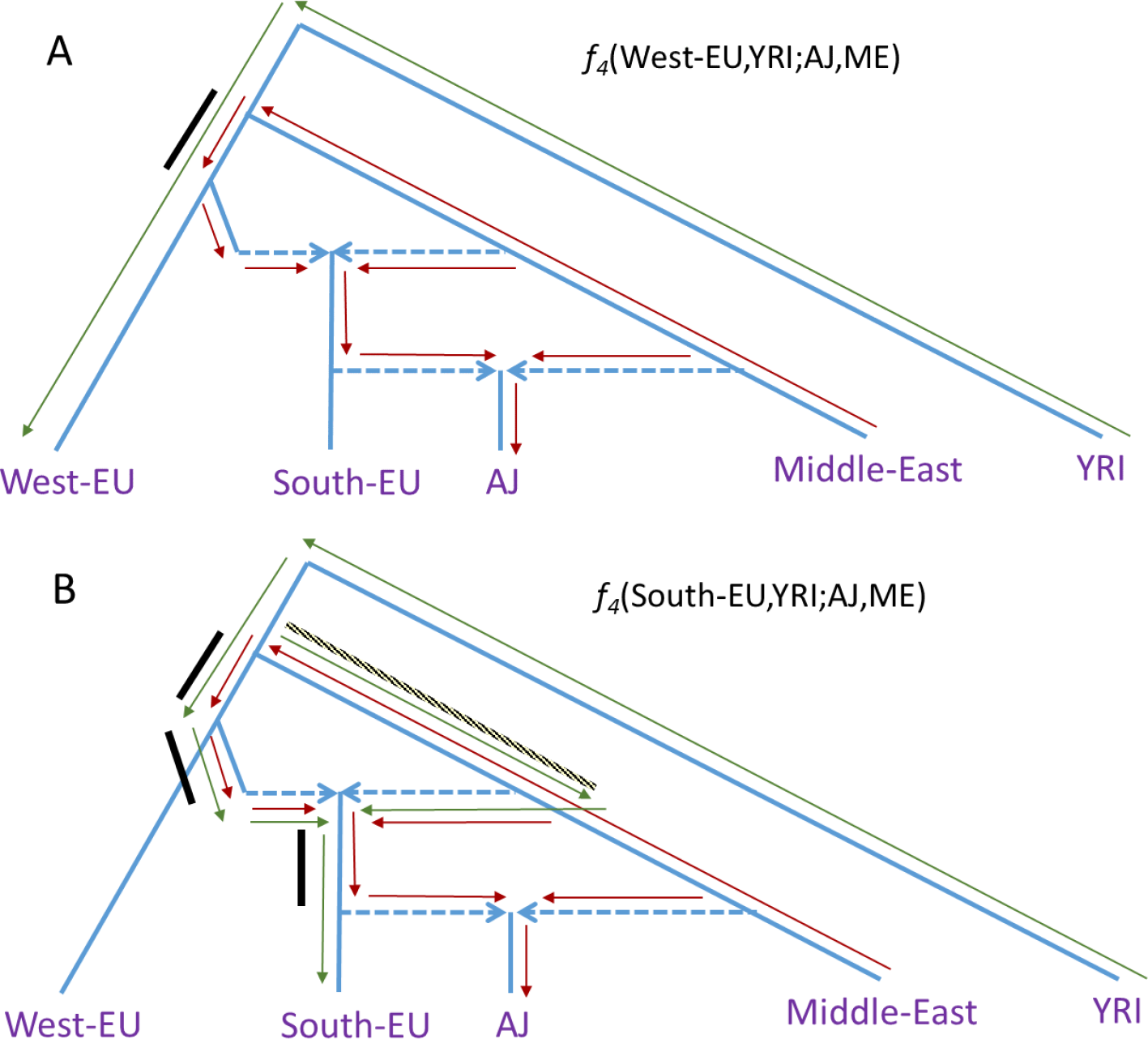
The effect of gene flow from the Middle East into Southern EU on *f_4_* statistics. Panels (A) and (B) demonstrate *f_4_*(West-EU,YRI;AJ,ME) and *f_4_*(South-EU,YRI;AJ,ME), respectively (cf Figure S4A). Lineages from the Middle-East into AJ are indicated with red arrows; lineages from YRI to Western or Southern Europe with green arrows. The *f_4_* statistic is proportional to the total overlap between these lineages (black bars). Whereas panel (B) *f_4_*(South-EU,YRI;AJ,ME)) has more overlapping branches than in (A), migration from the Middle-East into South-EU introduces a branch where the arrows run in opposite directions (patterned bar). Hence, the observed *f_4_* statistic in (B) may be lower (depending on the branch lengths) than in (A), even if Southern EU is the true source of gene flow into AJ.

**Figure S6.**
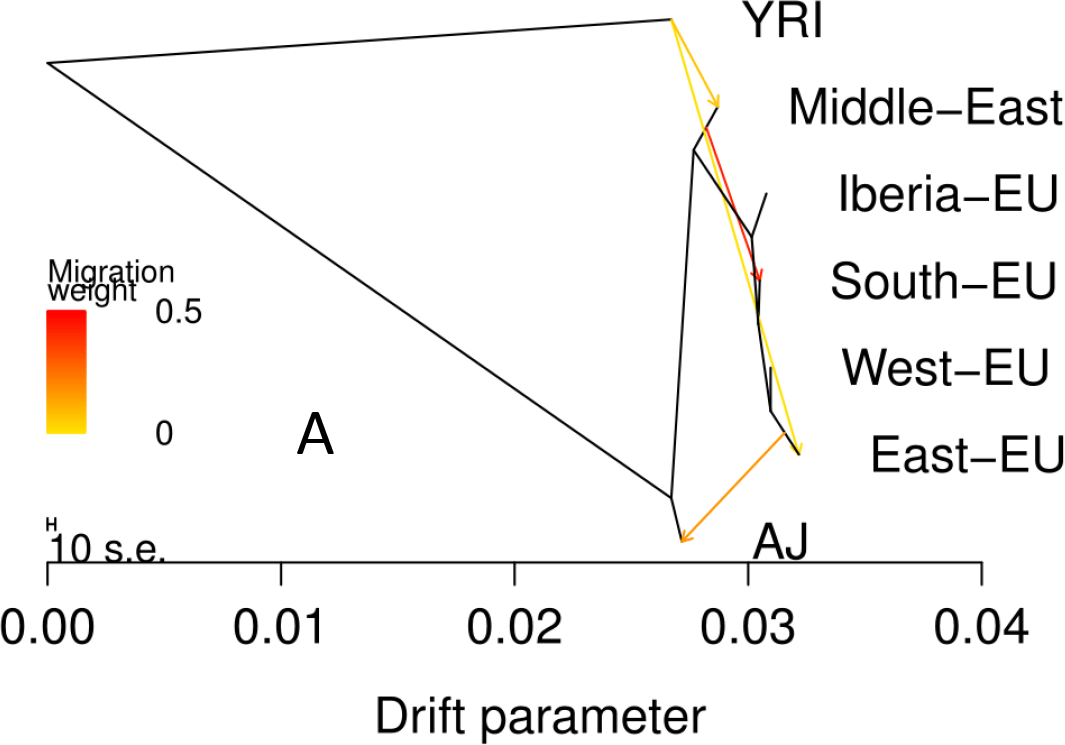
The graph structure of the AJ/EU/ME population histories, as estimated by *TreeMix* [50]. (A) Real data. (B) Simulated AJ data (along with the EU and ME populations in our study). Two hundred AJ genomes were simulated according to a 4-way model with 50% Middle-East, 35% South-EU, 12% East-EU, and 3% West-EU ancestries, with the mixing occurring 30 generations ago. The arrows indicate gene flow.

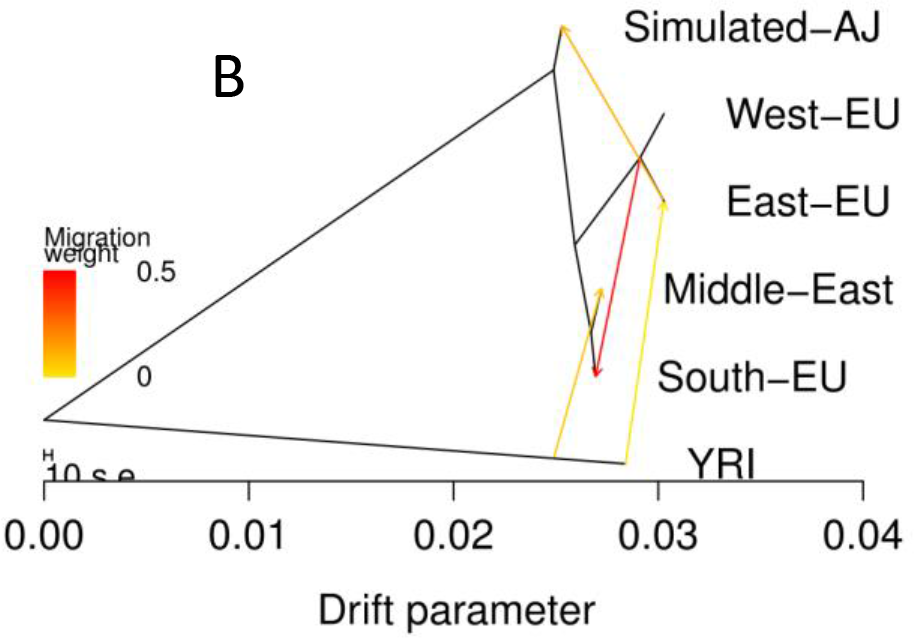

## Supplementary Text

### Supplementary Text S1: Local ancestry inference

#### Testing local ancestry inference (LAI) tools

LAI tools are known to be highly effective for distinguishing ancestries at a continental level (e.g., African vs European ancestry); however, at the subcontinental level, LAI might be noisy. Thus, before selecting an LAI tool, we used simulated admixed genomes from pairs of populations from the 1000 Genomes Project (phase I [1]) to determine the accuracy of *LAMP-LD* [2] and *RFMix* [3] (as reflected by the proportion of sites whose ancestry was correctly classified). Both programs use a window-based framework; *LAMP-LD* uses a generative approach using Hidden Markov Models, whereas *RFMix* uses a discriminative modeling approach via conditional random fields parameterized by random forests. For our initial tests, we used the 1000 Genomes Project data (phase I [1]) and found that while *LAMP-LD* was effective for distantly related populations (e.g., admixture between YRI and CEU), it had a much lower resolution for closer populations (e.g., TSI and FIN; populations with F_st_ around 1%, about the same as that between AJ and EU/ME populations). In contrast, *RFMix* was more effective at distinguishing TSI/FIN ancestries, and subsequent analyses demonstrated its capacity to distinguish (albeit with noise) also between Middle-Eastern and European ancestries. We therefore decided to use *RFMix* for all subsequent analyses.

#### Robustness to phasing errors

We note that while our local ancestry inference pipeline ran on perfectly phased data for our simulations, the AJ genotypes were only computationally phased. To determine whether phase switch errors are a concern, we performed the following experiment. We simulated 100 individuals with admixture occurring 30 generations ago and ancestry proportions 50% Southern European and 50% Levantine. After pairing sets of simulated chromosome, we randomly scrambled the phase, and then ran *Shapeit* to computationally re-phase all genotypes. We then re-ran the simulated genomes through our entire pipeline to infer the most likely geographic source. We found that the results essentially remained the same as when working with perfect phase, namely the genomes were localized to the true underlying European and Middle-Eastern subcontinental ancestry (Southern Europe and Levant) and the number of sites correctly classified as EU/ME did not change. Since computationally phasing each set of simulated genomes would have been extremely computationally expensive, the original phased simulated genotypes were used in all analyses.

#### The effect of filtering low-quality SNPs

We initially filtered out SNPs according to *RFMix’*s posterior probability (a measure of the confidence of the SNP coming from a specific ancestry), as we observed in simulations that filtering led to higher accuracy of LAI. However, we found that filtering led to biases in our geographic localization pipeline (based on the frequencies of the alleles at the EU segments). Specifically, we found that we were able to correctly localize a Southern European source only when we did not filter any SNPs. We attribute this result to the Middle-Eastern gene flow into Southern European (specifically, Italian) populations (e.g., [4]) and our use of a diverse reference panel that includes multiple European ancestries. These are expected to result in lower confidence in classifying Southern EU segments as European compared to segments from other European sources. In turn, filtering low quality SNPs would lead to disproportionately retaining segments of Northern European origin, thus wrongly localizing the EU segments even if the true source is Southern Europe. To guarantee the unbiased nature of our pipeline, we therefore did not filter any SNPs in all subsequent analyses.

### Supplementary Text S2: *PCAMask*

*PCAMask* is a software tool that performs principal component analysis restricted to the SNPs in each individual that derive from a specific ancestry [5, 6]. In theory, such a tool should be able to pinpoint the subcontinental ancestries of admixed individuals, but the utility of *PCAMask* on admixture between closely related populations was unknown. Running PCAMask on the AJ genomes (along with the reference panels described in the main text), we found that occasionally, the European component of the AJ genomes clustered around Southern Europe and that the Middle Eastern component of the AJ data clustered around the Levant region, in concordance with the results we present in the main text. Nevertheless, we did not include these results due to a number of technical issues (see also [7], which raised similar issues). Specifically, we found that in certain situations, the algorithm did not reach convergence and some AJ individuals were localized far away from the main AJ cluster. In addition, we found that the program did not appear to control for the number of admixed individuals: we noticed that increasing the number of AJ individuals led to their inconsistent placement. Finally, we compared the clustering of the reference EU and ME individuals between *PCAMask* and the commonly used *SmartPCA* tool [8], and noticed discrepancies in the clustering pattern. We therefore leave a more rigorous interpretation of *PCAMask’*s results to future work.

### Supplementary Text S3: GLOBETROTTER

#### Comparing EU ancestry proportion estimates between *RFMix* and *GLOBETROTTER*

The estimate of the total EU ancestry from the *RFMix* analysis came out as 53%, which is consistent with our previous estimate of ≈50-55% based on whole-genome data [9], as well as the estimate from the *f_4_* analysis (when calibrated by simulations). In contrast, the estimate from *GLOBETROTTER* [10] was 70% (among which 55% was Southern European). We find that reconciling these estimates is difficult, as evidence exists to support both the LAI-based estimate and the *GLOBETROTTER* based estimate.

To test *GLOBETROTTER*, we simulated individuals with ancestry proportions 8% Western EU, 8% Eastern EU, 34% Southern EU, and 50% Levant, which all admixed 30 generations ago. *GLOBETROTTER* was able to recover all proportions within α1% of the simulated ones. For simulations with ancestry proportions 70% Southern EU and 30% Levant, the *GLOBETROTTER* inferred EU proportions were slightly overestimated at 73%, implying 67% EU ancestry in AJ. On the other hand, the *RFMix* inferred proportions were underestimated at 62%. However, the bias for simulated 50% Southern EU and 50% Levant ancestries was lower, with *RFMix* inferred EU proportions at 48%.

Additional support to the *RFMix* estimate came from simulations of admixture 30 generations ago, with proportions 8% Western EU, 8% Eastern EU, Southern EU proportions varying between 20% to 80%, and the remaining proportions from the Levant. We then applied the geographic localization pipeline described in the main text, and compared the number of chromosomes having the maximum likelihood at Southern Europe. The best match to the real data was obtained when simulating 35% Southern EU ancestry (leaving 49% Levantine ancestry), in agreement with the direct estimate.

In conclusion, there remains some uncertainty regarding the amount of EU ancestry in AJ, to be fully resolved in future studies. It seems plausible that the true EU ancestry proportions are around ≈60%, midway between the *RFMix* and the *GLOBETROTTER* estimates. For most of this paper we assumed the *RFMix* estimate (≈55%), as (i), it is supported by other lines of evidence; (*ii*) the results from the two modes of *GLOBETROTTER* were discordant. *GLOBETROTTER*’s ancestry profiles were obtained for each AJ chromosome independently, and thus should not be confounded by the severe AJ bottleneck in an obvious way [11]; however, more subtle confounding is possible.

#### *GLOBETROTTER*-inferred admixture parameters on simulated data

We used simulations to test the ability of *GLOBETROTTER* to infer admixture time and sources [10]. The simulated individuals had 70% Southern EU and 30% Levant ancestries, with admixture occurring 30 generations ago. *GLOBETROTTER* inferred two sources: the first, comprising of 39% of the total ancestry, was a mixture of 15% Southern European ancestry and 85% Levant ancestry; the second source was 1% Eastern European, 28% Western European, and 71% Southern European. Thus, the true Southern EU ancestry proportions were not properly recovered (inferred 49% vs simulated 70%), although the global EU ancestry was inferred correctly (67% vs simulated 70%). The inferred admixture time was overestimated at 40 generations.

#### The number of admixture events

*GLOBETROTTER* is able to infer multiple admixture events, although for AJ, the inferred history included only a single event. This might be at odds with our hypothesis (supported by the IBD analysis) of pre-bottleneck admixture with Southern Europeans followed by post-bottleneck admixture with (possibly) Eastern Europeans. However, we note that one source of ancestral population inferred by GLOBETROTTER is a mixture of Southern EU and Levant, which may correspond to the earlier event. Additionally, the two events may be too close together to be teased apart, and the inference of admixture times might be confounded by the severe AJ bottleneck [10].

## Supplementary Text S4: The distribution of ancestry proportions under two-wave admixture

### 1 The distribution of ancestry proportions under general distributions of segment lengths

In the main text, we considered a simple admixture pulse model, under which the distribution of segment lengths in **A** and **B** is exponential with rates (1 − *m*)*t* and *mt*, respectively. Under this model, the distribution of ancestry proportions was available in a closed form. Under a more complex admixture history, we assume that the distribution of the length of **A** and **B** segments take the general form *q*A(*ℓ*) and *q*_B_(*ℓ*). We still assume that **A** and **B** segments are independent (see below). The process can then be modeled as a two-state process. We start on the left end of the chromosome in state **A** or **B** with probabilities *p_A_* = 〈*ℓ_A_*〉 /〈*ℓ_A_*〉 +〈*ℓ_B_*〉) and 1 − *p_A_*, respectively (where 〈*ℓ_A_*〉 and 〈*ℓ_B_*〉 are the mean segment lengths), and draw a random segment length from the selected ancestry. When the first segment terminates, we switch ancestries and draw a segment length from the other ancestry, and so on until we reach the end of the chromosome.

The distribution of *x*, the **A** ancestry proportion, can be computed in Laplace space by extending renewal theory methods developed in the physics domain (e.g., [1, 2]). Let s be the Laplace pair of *L* (the total chromosome length) and *u* as the Laplace pair of *L_A_* = *xL* (the total chromosome length covered by A segments). We then transform the density *f*(*L_A_; L*) (from which the density of *x* can be easily obtained) to 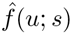. After some calculations using renewal theory, we eventually obtain,

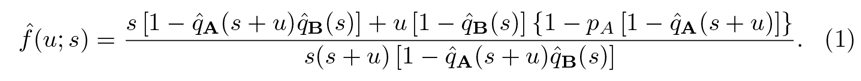

In the above equation, 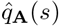 and 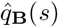 are the Laplace transform (*ℓ* → *s*) of *q*_A_(*ℓ*) and *q*_B_(*ℓ*). The details of the derivation are somewhat tedious and are therefore omitted. It can be shown, using Eq. (1), that the mean ancestry proportion 〈*x*〉 approaches p_A_ as *L* → ∞ It can be also shown that Eq. (1) reduces to Eq. (1) in the main text for the admixture pulse model.

### 2 Conditions under which consecutive segments are independent

To study complex admixture histories, we use the model developed by Gravel [3] (section *General incoming migration in the absence of drift* and Figure 3 there). Gravel proposed that the ancestry along the chromosome could be described by a Markov process, whose states correspond to the identity of the source population (i.e., **A** or **B**), combined with the time when each segment entered the admixed population. Gravel then derived the transition rates for any general admixture history. While the extended state space process is Markovian under any history, consecutive **A** and **B** segment lengths are no longer independent. However, further examination demonstrates that as long as migration beyond the the initial event is limited to just one population, consecutive segment lengths remain independent.

### 3 A two-wave admixture model

Consider a model where populations **A** and **B** have merged *t*_1_ generations ago, contributing proportions *m* and 1 − *m* to the admixed population. Then, *t*_2_ (*<t_1_*) generations ago, migrants from population **A** have replaced a proportion *μ* of the gene pool of the admixed population. No other events then take place until the present. The corresponding Markov process, using Gravel’s method [3], has three states: *A*_1_, *A*_2_, and *B*, representing migrant segments from **A** at time *t*_1_, from **A** at time *t*_2_, and from **B** (at time *t*_1_), respectively. Let us compute the distributions of the lengths of **A** and **B** segments.

The transition rate is *t*_1_ when at states *A*_1_ and *B*, and *t*_2_ when at *A*_2_. It can be shown that once a transition is made, the next state is chosen according to the following transition probability matrix

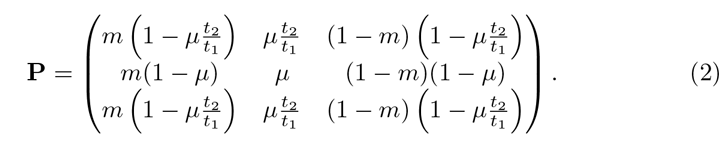

The states are ordered as (*A*_1_,*A*_2_,*B*) and **P***_ij_* (*i,j* = 1, 2, 3) is the probability to jump from state *i* to state *j*. Note that we neglected the first generation after admixture, during which **A** and **B** segments do not yet mix [3].

It is now easy to see that **B** segment lengths are distributed exponentially with rate *t*_1_(1 − **P***_B,B_*), or

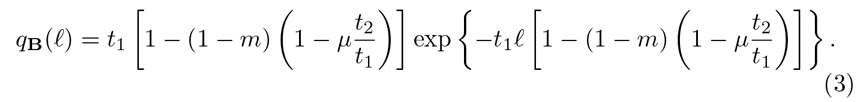

This equation was also (implicitly) derived in [4] in a different way. For the **A** segments, define *q_A_1__* (*ℓ*) as the distribution of **A** segment lengths, *when the process entered any of the **A** states at state A_1_*, and similarly for *q_A_2__* (*ℓ*). Since the process enters *A_1_* and *A_2_* from *B* (with the possible exception at the leftmost end of the chromosome), the distribution of **A** segments therefore satisfies

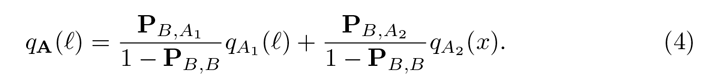

To find *q_A_1__* (*ℓ*) and *q_A_2__* (*ℓ*), ^we can^ write integral equations,

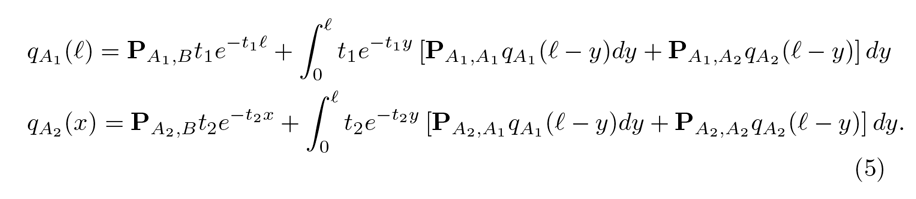

We solved those equations by Laplace transforming them (*ℓ* → *s*). Using the convolution theorem,

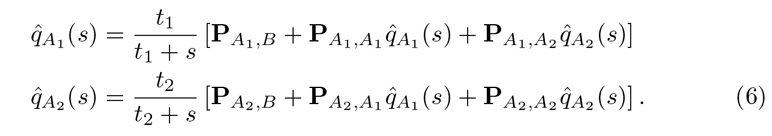

These are two linear equations in two variables (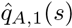 and 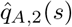 that are easily solved. Then, *q_A,l_*(*ℓ*) and *q_A,2_*(*ℓ*) be obtained by Laplace transform inversion. We then use Eq. (4) to obtain q_**A**_(*ℓ*). We carried out these steps in Mathematica, leading to the final result,

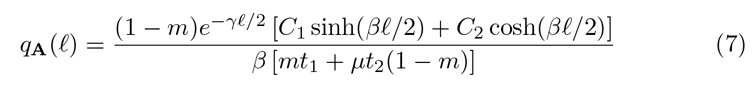

where 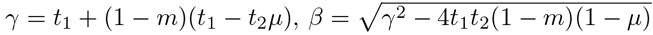, 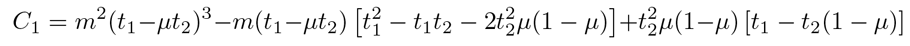 and

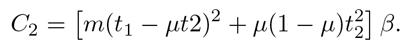

Now that we have *q_A_* and *q_B_* (Eqs. (7) and (3), respectively), we can use Eq. (1) for the distribution of the ancestry proportions. We inverted 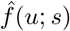 with respect to *u* using Mathematica and then numerically with respect to *s* to obtain *f*(*x; L*).

We note that we can also view the second migration wave as gene flow coming from a *third* population. Our results here and in the main text then automatically provide the distribution of ancestry proportions coming from each of the three sources.

### 4 Simulation results and fitting

We ran simulations of the Markovian Wright-Fisher model described by Gravel [3]. The model assumes 2*N* haploid individuals (chromosomes). Each chromosome in the current generation is formed as a mixture of the chromosomes of the previous generation. Ancestry changes occur as a Poisson process with rate 1 (Morgan), and at each ancestry change, the ancestral chromosome is chosen randomly out of all 2*N* available chromosomes. In the pulse admixture model, each chromosome in the first generation is assigned to population **A** or **B** with probabilities m and 1 − *m*, respectively, and the evolution of the chromosomes is traced for *t* generations. The two-wave model is the same (with overall time *t*_1_), except that at *t*_2_ generations ago, each chromosome is replaced by a whole-**A** chromosome with probability *μ*.

Representative simulation results are shown in Supplementary Text Figure 1. It can be seen that our theory matches the empirical data very well. However, the empirical distribution can also be fitted very well by a distribution corresponding to an admixture pulse model, with parameter *m*_pulse_ close to the expected mean (*μ* + *m* (1 − *μ*)) and *t*_pulse_ intermediate between *t*_1_ and *t*_2_. This suggests that almost any inference based on the more complex model will not have sufficient evidence to justify the additional admixture event.

**Supplementary Text Figure 1:**
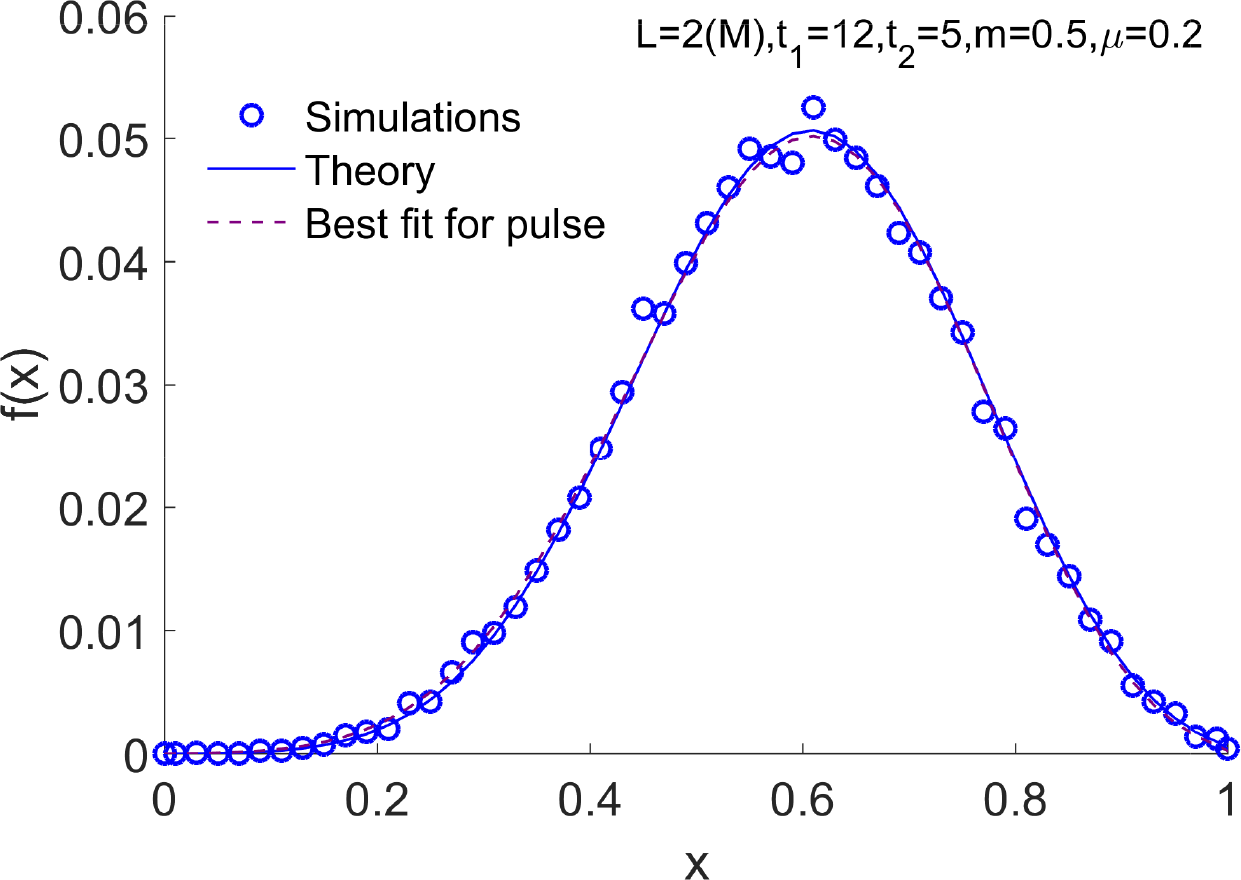
Two-wave admixture: simulations and theory. We simulated a two-wave admixture model according to a Markovian Wright-Fisher model [3] with *N* = 2500. The other model parameters are indicated on top of the figure. We recorded the fraction of each chromosome that descends from the **A** population, and plotted the histogram of the ancestry proportions (circles). The theory that we developed (Eqs. (1), (3), and (7)) is plotted as a solid (blue) line. We then fitted a pulse admixture model with just two parameters (*m* and *t*) by matching the mean and variance of the empirical data. The distribution of the ancestry proportions under the pulse model (Eq. (1) in the main text) is plotted as a dashed (purple) line. The best fit for *t* was 9.7, intermediate between *t*_1_ and *t*_2_.

